# Diurnal transcript profiling of the diatom *Seminavis robusta* reveals adaptations to a benthic lifestyle

**DOI:** 10.1101/2020.11.23.393678

**Authors:** Gust Bilcke, Cristina Maria Osuna-Cruz, Marta Santana Silva, Nicole Poulsen, Sofie D’hondt, Petra Bulankova, Wim Vyverman, Lieven De Veylder, Klaas Vandepoele

## Abstract

Coastal regions contribute an estimated 20% of annual gross primary production in the oceans, despite occupying only 0.03% of their surface area. Diatoms frequently dominate coastal sediments, where they experience large variations in light regime resulting from the interplay of diurnal and tidal cycles. Here, we report on an extensive diurnal transcript profiling experiment of the motile benthic diatom *Seminavis robusta*. Nearly 90% (23,328) of expressed protein-coding genes and 66.9% (1124) of expressed long intergenic non-coding RNAs (lincRNAs) showed significant expression oscillations and are predominantly phasing at night with a periodicity of 24h. Phylostratigraphic analysis found that rhythmic genes are enriched in deeply conserved genes, while diatom-specific genes are predominantly associated with midnight expression. Integration of genetic and physiological cell cycle markers with silica depletion data revealed potential new silica cell wall associated gene families specific to diatoms. Additionally, we observed 1752 genes with a remarkable semidiurnal (12-h) periodicity, while the expansion of putative circadian transcription factors may reflect adaptations to cope with highly unpredictable external conditions. Taken together, our results provide new insights into the adaptations of diatoms to the benthic environment and serve as a valuable resource for diurnal regulation in photosynthetic eukaryotes.

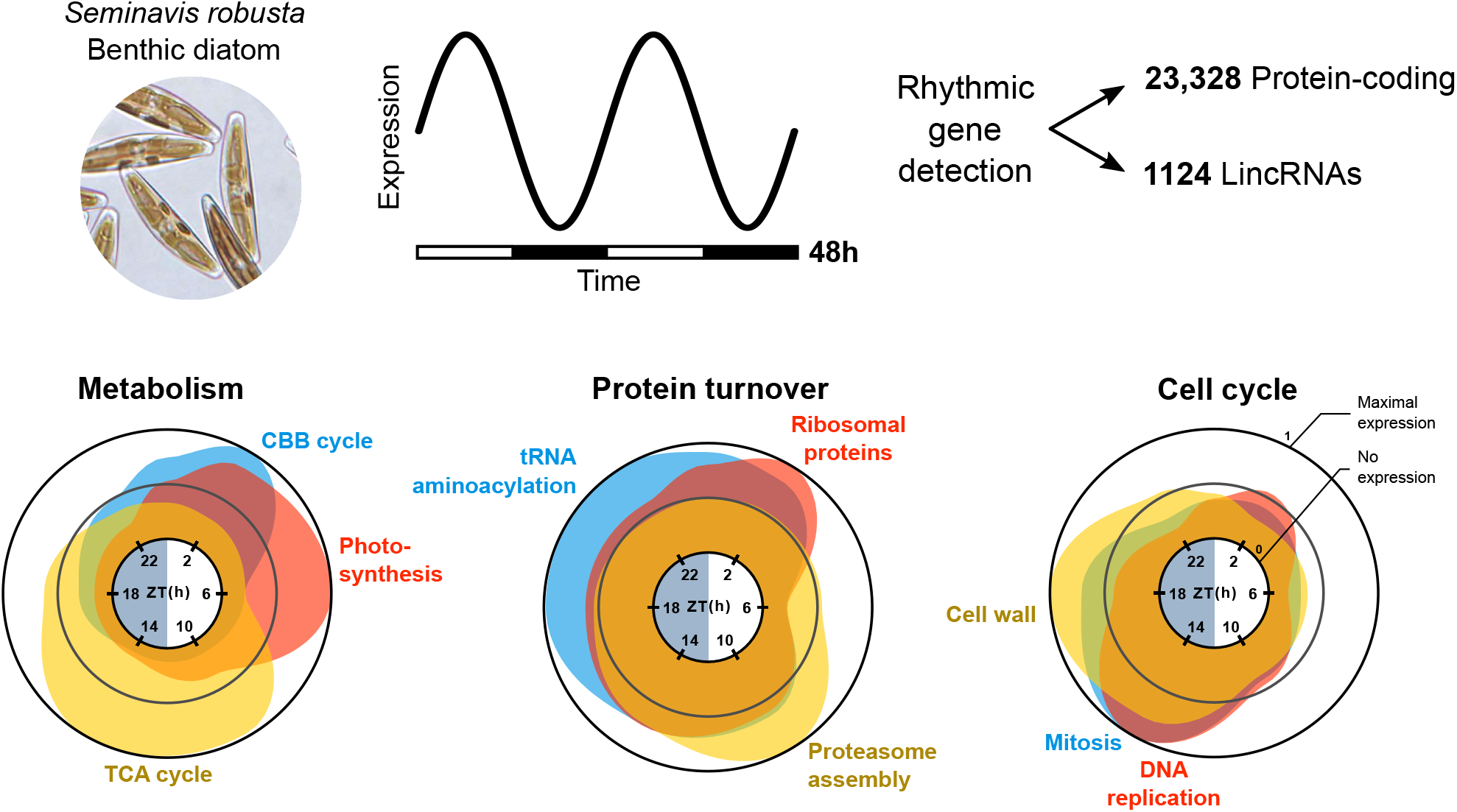

## 2 Introduction

Most organisms are exposed to a daily alternation of light and darkness as the result of the earth’s rotation. This has led to the evolution of rhythmic cellular programs that are controlled by environmental cues as well as endogenous circadian clocks [1]. In particular, coordination with diurnal cycles (day/night cycles) is crucial for photosynthetic organisms, whose metabolism relies heavily on the availability of sunlight [2]. Compared to multicellular plants, rhythmicity is especially pronounced in microalgae, where up to 80% of the transcriptome is subject to rhythmic oscillations [3, 4].

Diatoms are prominent stramenopile microalgae in marine and freshwater environments. They account for up to 20% of annual global primary production and play an important role in biogeochemical cycling of nutrients in aquatic ecosystems, including silica [5]. Diatom cells are characterized by a unique bipartite silica cell wall, called the frustule. Although cell wall structure and morphology is widely used in diatom taxonomy and biodiversity studies, silica biomineralization is still poorly understood [6].

To date, only a handful of studies have explored rhythmic gene expression in diatoms, generally with a particular focus on the response of the diurnal transcriptome to nutrient limitation [7, 8, 9, 10]. The most extensive accounts come from the pennate diatom *Phaeodactylum tricornutum*, whose diel coordination of carbohydrate and cell cycle pathways was shown to be strongly affected by iron availability and for which the first diatom circadian regulator was recently characterized [9, 10, 11]. Furthermore, studies thus far have focused solely on planktonic diatom species, which typically inhabit well-mixed water columns. In contrast, no data are available on diurnal variation in gene expression of the extraordinarily diverse group of benthic pennate diatoms, dominating some of the most productive ecosystems on the globe [12, 13, 14]. Yet, benthic ecosystems are exposed to dynamic and often extreme light regimes, resulting from the combined effects of tidal phasing, sediment composition and water column turbidity [15, 16, 17]. Under laboratory conditions, benthic diatom species generally show maximal growth rates at irradiances of 10-50 μE m^−2^ s^−1^, similar to conditions of shading that are periodically encountered in benthic communities [18, 19, 20, 21, 22]. Not only light availability varies over the course of the diel cycle, also nutrient and oxygen availability in the sediment show rhythmic patterns [23, 24], adding to the complexity and dynamic nature of benthic habitats. Motile species from the raphid pennate diatom clade are particularly successful in these environments and use the secretion of mucilaginous exopolysaccharides (EPS) to actively migrate through the sediment at dusk and dawn to maximize photosynthesis while avoiding excessive grazing [24, 25, 26, 27]. This suggests that benthic diatoms employ elaborate mechanisms to track the diurnal cycle in order to optimize photosynthesis, nutrient uptake and survival.

In recent years, *Seminavis robusta* has gained interest as a model species for benthic diatoms and to study raphid pennate life cycle regulation. This relatively large-sized (~70 μm initial cell size) raphid pennate diatom inhabits shallow marine and brackish zones and has been reported from the North Sea, the Mediterranean and the Pacific ocean [28, 29, 30]. In culture, *S. robusta* cells display typical benthic behaviour, including surface attachment, biofilm formation and vigorous gliding motility, which has been exploited to study chemotaxis [31]. Cultures from strain 85A have been popular for transcriptomics experiments [32, 33, 34, 35]. This strain is a fifth generation lineage from the *S. robusta* laboratory pedigree whose ancestral clones originate from subtidal sediments of the Veerse Meer in The Netherlands (51°32’36” N, 3°48’15” E) [36, 37]. Many aspects of the physiology of *S. robusta* appear to be tied to light availability and diurnal cycles. Prolonged darkness followed by illumination causes synchronization in cell cycle progression, chloroplast repositioning, pigment synthesis and sexual reproduction [38, 39]. Furthermore, *S. robusta* responds to high irradiances both by downwards migration into the sediment and the expression of photoprotective LHCX genes [40, 41]. The recent generation of a reference genome assembly and the construction of a transcriptome atlas that spans various environmental conditions, biotic interactions and life cycle stages represents an important resource for comparative genomic studies, allowing to identify genomic features associated with the lifestyle of obligate benthic diatoms [33].

Long non-coding RNAs (lncRNAs) are an emerging class of transcripts longer than 200 nucleotides that are not translated into protein [42]. Advances in whole-transcriptome sequencing have revealed that a considerable portion of the non-coding DNA in plant and animal genomes make up lncRNAs, which are expressed at low and mostly tissue-specific levels [42, 43]. Despite lncRNAs still being largely unexplored, they appear to play a pivotal role in gene expression regulation and are increasingly implicated in various biological processes such as environmental stress and defense [44]. In addition, numerous lncRNAs are involved in diel regulation or the circadian clock in both plants and animals [45, 46]. Although intergenic lncRNAs (lincRNAs) have been described in diatoms in the context of environmental stimuli [47, 48], rhythmic oscillations in these genes have not been assessed yet.

In this study, we investigated the diurnal regulation of the benthic diatom *S. robusta* using RNA-sequencing (RNA-seq). Sampling of a two-day (48h) time series on three separate occasions allowed us to identify genes with a stable recurring expression oscillation. Next to the analysis of protein-coding genes, lincRNAs were annotated in the *S. robusta* genome to identify the first rhythmic lincRNAs in microalgae. We further performed phylogenomic and comparative transcriptomic analyses to answer outstanding questions regarding the conservation of rhythmic gene expression in benthic versus planktonic diatoms, the evolutionary history of rhythmic gene phasing and periodicity, and synchronization of organelle division to the diurnal cycle. Finally, we took advantage of available silicon depletion transcriptome datasets to uncover several new gene families with a potential role in diatom cell wall synthesis.

## 3 Results and discussion

### 3.1 The majority of *Seminavis robusta* genes exhibit a rhythmic expression pattern under diel growth conditions

We maintained *S. robusta* cultures in a 12h-light/12h-dark photoperiod to identify genes exhibiting periodic expression. Three replicate cultures were sampled over a 48-h time course at 4-h intervals beginning at ZT2 (Zeitgeber Time = hours since the onset of illumination), resulting in 12 time points over two complete diel cycles (Fig. 1a). After RNA sequencing, more than one billion reads were obtained with an average of 28 million reads per sample and an average mapping rate of 87.15%. Multidimensional scaling revealed a concerted diel progression in gene expression, with samples harvested at the same time of each day clustering together, regardless of the experimental repeat (Fig. 1b). Next, a rhythmicity-detection algorithm was used to derive genes with a significantly cyclic expression pattern, and retrieve the periodicity and phasing of each rhythmic gene’s waveform (Hutchison et al., 2015). Genes having significant oscillatory gene expression (q-value < 0.05) were classified into clusters by period (24h or 12h) and phase (ZT2, ZT6, ZT10, ZT14, ZT18 and ZT22) (Fig. 1c,d, Suppl. Fig. 1-2). Additionally, we partitioned 24-h periodic rhythmic genes across more general categories based on their maximum and minimum expression: day (ZT2, ZT6, ZT10), night (ZT14, ZT18, ZT22), dawn (ZT22 or ZT2) and dusk (ZT10 or ZT14) (Fig. 1a,c). These hierarchical gene sets (rhythmicity > periodicity > phasing) were subsequently used to assess the rhythmic properties of biological processes and the evolutionary conservation of rhythmicity.

**Figure 1:**
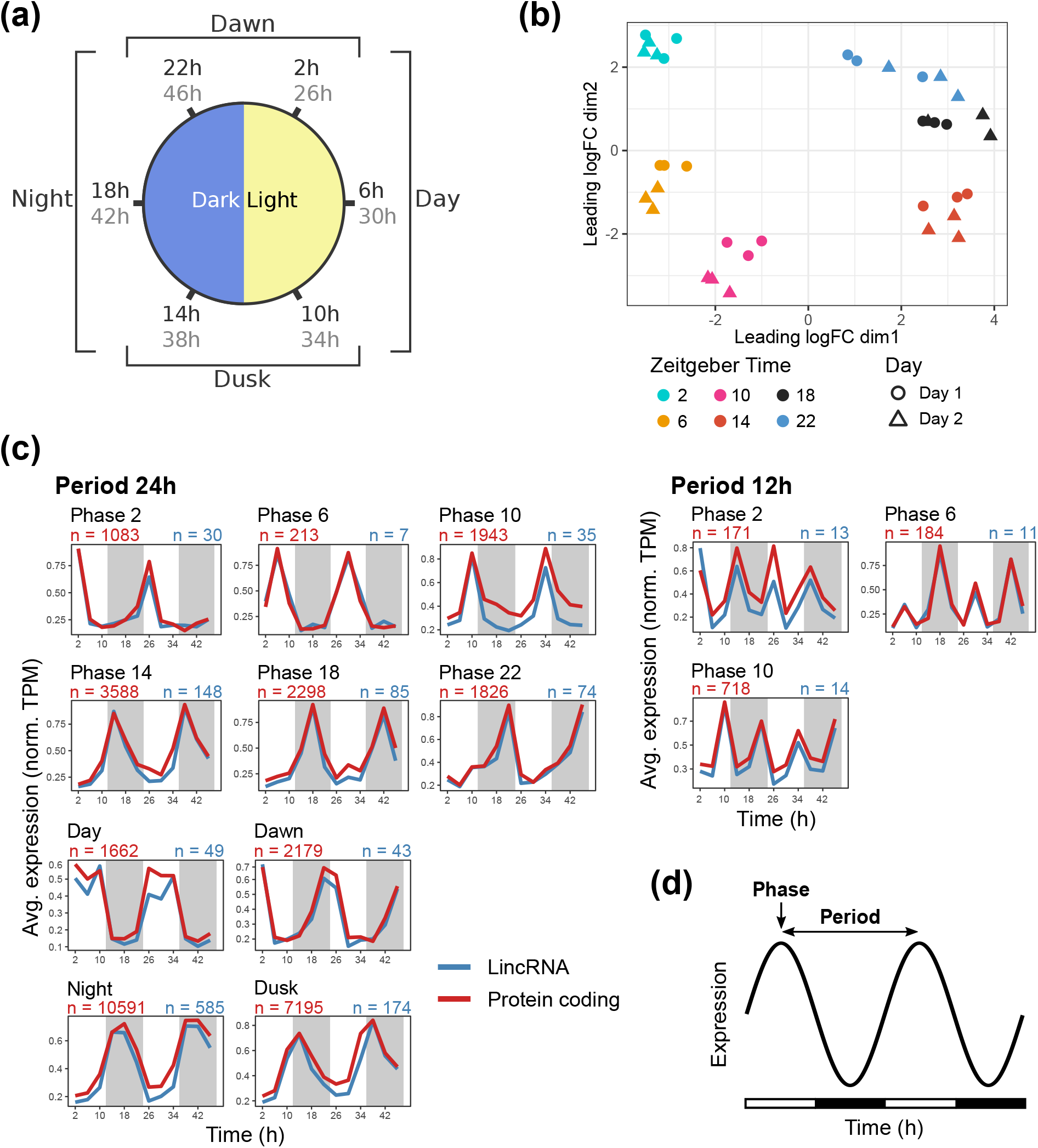
Overview of the diurnal gene expression experiment and classification of rhythmic genes in *S. robusta*. **(a)** Schematic representation of the experimental setup and classification. The blue and yellow shaded semicircles represent the dark and light part of the 12h/12h cycle, respectively. Samples acquired in the first 24h are printed in black, while the corresponding samples from 24h-48h are labelled in grey. Genes were classified in clusters for each phase (peak expression) as well as broader categories “Day”, “Night”, “Dawn” and “Dusk”. **(b)** Multidimensional scaling (MDS) plot of 36 sequenced samples for the top 500 most differential genes of the diurnal time series. Colours indicate time of day (Zeitgeber Time, hours since first illumination) and shapes represent whether a sample belongs to the first or second day of the experiment. **(c)** Average expression for 24-h and 12-h period genes, organized in temporal classes by their predicted phase. Expression in transcripts per million (TPM) was normalized to a maximum of 1. “n” represents the number of genes in each cluster, printed in red and blue for protein-coding genes and lincRNAs, respectively. **(d)** Nomenclature of rhythmic expression waveforms. The fitted expression of a hypothetical gene during the time series is presented, with day and night treatment indicated in white and black bars. Terms used throughout the paper to discuss properties of the waveform are indicated with arrows.

A total of 23,328 protein-coding genes showed a significant rhythmic expression pattern. This corresponds to 89% of protein-coding genes that were expressed in our diurnal dataset, and 64% of all protein-coding genes encoded in the *S. robusta* genome (Suppl. Table 1). Proportions in this range have been described for various unicellular algae [3, 4, 49, 50]. Lower levels of rhythmicity were reported in the planktonic diatoms *P. tricornutum* and *T. pseudonana*, although this could be due to differences in sample resolution and rhythmicity detection algorithms [7, 8, 9]. Re-analysis of a comparable, 24-h *P. tricornutum* RNA-seq dataset confirmed a relatively low proportion of rhythmic genes compared to *S. robusta*, assessed by the same algorithm (Suppl. Fig. 3, Suppl. Table 1) [10]. The higher level of rhythmicity of *S. robusta*’s transcriptome might be explained by the longer duration of the time series or experimental confounders such as light intensity and cell density. Alternatively, it could reflect an adaptation of benthic diatoms to cyclic changes in the abiotic conditions of their environment [15, 51, 23, 24]. These pronounced transcriptomic oscillations occurred under an experimental irradiance of 30 μE m^−2^ s^−1^, suggesting that diurnal coordination is possible even under conditions of shading by the overlying water column [22, 16].

Most rhythmic protein-coding genes displayed a 24-h (92.5%) or 12-h (7.5%) periodicity, and similar to reports in other diatoms, were predominantly phasing at dusk and at night (Fig. 1c) [10, 7, 8]. We cross-referenced the 10,145 protein-coding genes that were not expressed during the diurnal cycle in a publicly available *S. robusta* expression atlas that reports on transcriptional activity during sexual reproduction, abiotic stress and diatom-bacterial interactions (Suppl. Table 2) [33]. More than half (5,315) of non-diurnal genes were transcriptionally active in at least one condition, although at lower expression levels than diurnally expressed genes (Suppl. Fig. 4a). Calculation of the Tau index as a measure of condition specificity [52] showed that non-diurnal genes were expressed in a more condition-specific manner than diurnally expressed genes (average Tau 0.82 and 0.49 respectively, Wilcoxon rank-sum test p < 2.2e-16) (Suppl. Fig. 4b). Hence, while most diurnally expressed genes display a broad expression pattern not associated with a specific condition, genes that were not expressed during our diurnal experiment seem to be relevant under specific conditions such as sexual reproduction (Suppl. Fig. 4c).

### 3.2 Detection of rhythmic lincRNAs suggests the non-coding transcriptome is involved in diurnal responses

We next investigated the diel expression of 7,117 high-confidence lincRNAs that we annotated in the *S. robusta* genome leveraging different RNA-seq datasets (see Methods). The number of *S. robusta* lincRNAs is in line with the diversity of lincRNAs in the brown alga *Saccharina japonica* [53], but exceeds that in the diatom *P. tricornutum* by about 5-fold [47]. In the *S. robusta* expression atlas, lincRNAs were transcribed at lower levels than protein-coding genes (average maximal log2(TPM) of 3.0 and 5.1 respectively, Wilcoxon rank-sum test p < 2.2e-16, Suppl. Fig. 4a) and showed higher condition specificity, as expected (Average Tau 0.85 and 0.54 respectively, Wilcoxon rank-sum test p < 2.2e-16, Suppl. Fig. 4b). From a total of 1,680 diurnally expressed lincRNAs, 1,124 showed significant oscillations (66.9%, Fig. 1c). Similar to the protein-coding genes, rhythmic lincRNAs displayed repetitive peaks in expression associated with a specific time of the day and were predominantly phasing at night and at dusk (Fig. 1c). Applying the guilt-by-association principle using co-expression networks and functional information of protein-coding genes, we could attribute a biological process to 250 rhythmic lincRNA genes, implying a cooperation between protein-coding and lincRNA genes [54]. Rhythmic lincRNAs were predicted to function in photosynthesis, carbohydrate metabolism, translation, proteolysis, cell cycle, regulation of transcription and intracellular signal transduction. In contrast to protein-coding genes, the majority of all lincRNAs (5,437, 76.4%) was not diurnally expressed. Thus, although a considerable set of lincRNAs plays a role in diurnal conditions, the majority of lincRNAs is rather associated with specific stress environments, as previously reported in diatoms (Suppl. Fig. 4c) [47, 48]. Finally, only 103 lincRNAs displayed conservation in at least one other diatom species, of which 19 were significantly rhythmic. These results suggest a poor conservation of rhythmic lincRNAs between distantly related diatom species or a strong divergence in primary nucleotide sequence, as is widely observed for lncRNAs, interfering with reliable homology detection [55].

### 3.3 Evolutionary analysis of rhythmic genes points towards ancient daytime functions and diatom-specific expression at night

Phylostratigraphic analysis, which explores protein homology in a wide set of species to date the origin of specific genes, revealed that rhythmic protein-coding genes were enriched in eukaryote and stramenopile age, while non-significantly rhythmic genes displayed enrichment in species-specific genes (Fig. 2a). The latter group of genes was mainly enriched in genes upregulated under different stress conditions in the *S. robusta* expression atlas (Suppl. Table 2) [33], while the set of rhythmic genes was predominantly enriched in downregulated genes under these conditions (Fig. 2b).

**Figure 2:**
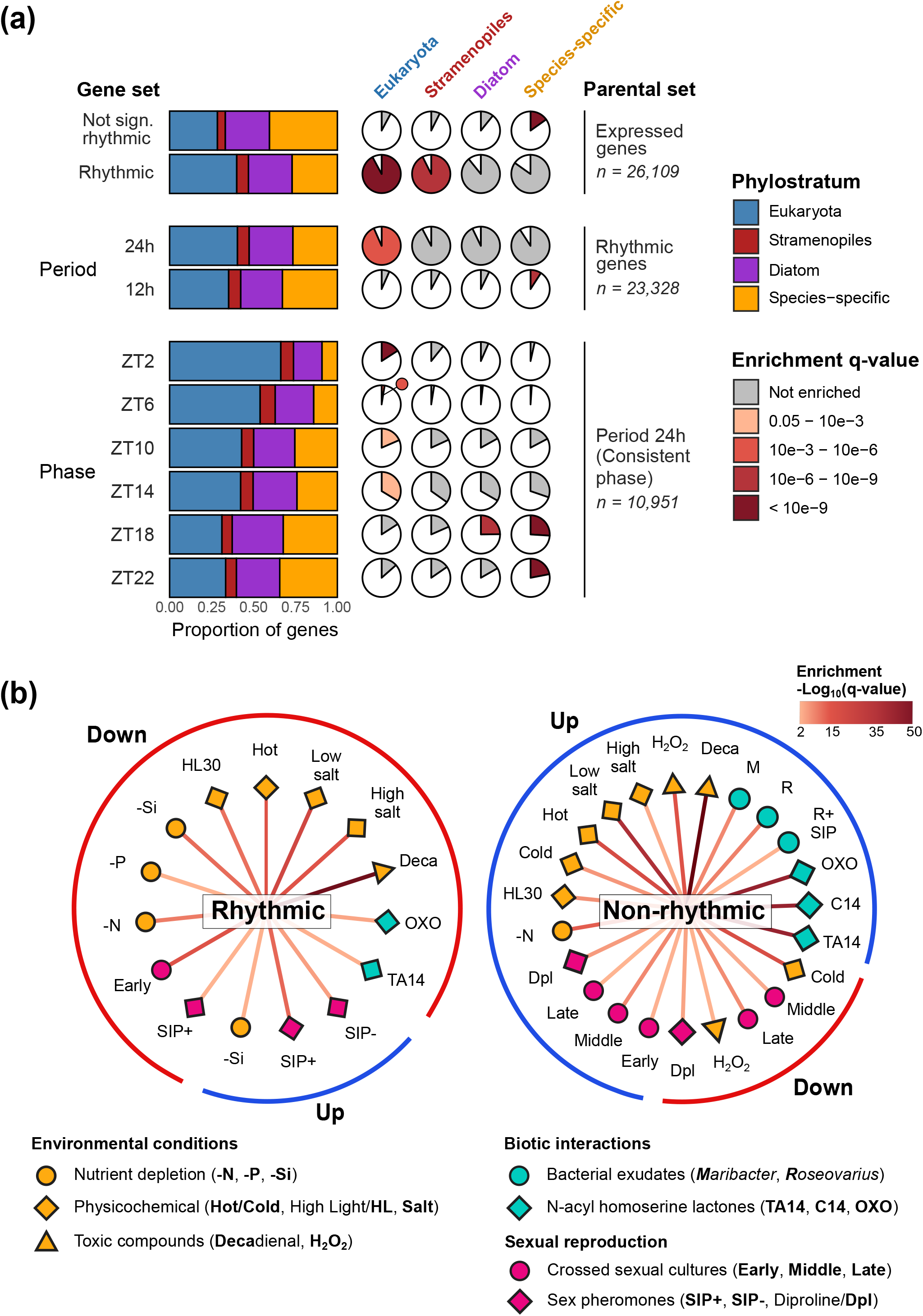
Phylostratigraphy and differential expression of rhythmic and non-significantly rhythmic protein-coding genes. **(a)** Distribution of gene ages within gene sets classified by rhythmic properties. Gene sets were defined at three hierarchical levels: rhythmicity, periodicity and phasing. The higher-level classification that gene sets belong to is indicated by “Parental Set” (n = number of genes in each parental set). For each *S. robusta* gene, a phylostratum (age class) was determined that ranges from “Eukaryota” (gene family occurs outside stramenopiles) to “species-specific” (gene family is restricted to *S. robusta*). The prevalence of phylostrata within gene sets is shown in two ways. (1) The proportion of each phylostratum in a gene set is displayed by a stacked bar plot. (2) Pie charts show the proportion of genes in a specific phylostratum (column, sums to 100%) that are classified into each gene set (rows). Enrichment of phylostrata for each gene set is displayed by the colour of the shaded areas, which relates to the q-value of enrichment. For example, rhythmic genes are significantly enriched for older phylostrata (Eukaryota and stramenopiles). This is reflected in the size of the pie charts: 92.2% and 92.6% of eukaryotic and stramenopile-age genes were rhythmic, while only 84.8% of species-specific genes were classified as rhythmic. **(b)** Network showing enrichment of differentially expressed genes over a wide range of conditions in the set of rhythmic (left) versus non-significantly rhythmic genes (right). The colour and shape of icons indicate classes of conditions included in the *S. robusta* expression atlas [33]. The text adjacent to each shape refers to the specific treatment, as detailed in bold in the legend at the bottom. -N, -P and -Si denote Nitrogen, Phosphorous and Silicon depletion respectively, and SIP stands for 3h of treatment with sex inducing pheromone. TA14, C14 and OXO14 are (derivatives of) N-acyl homoserine lactone quorum sensing compounds emitted by bacteria. More details about treatments of the *S. robusta* expression atlas can be retrieved from Suppl. Table 2. The significance of the enrichment for each condition is highlighted by a colour gradient from light red to dark red in −log10(q-value) scale in the edge of the network. The lines in the outer circle summarize the proportion of conditions enriched in down- (red) and upregulated (blue) genes.

These results suggest that phylogenetically old (conserved) rhythmic genes play a role in basal functions that are suppressed during external stress, such as cell cycle, photosynthesis and storage carbohydrate synthesis. Meanwhile, young and non-rhythmic genes are more important in specific environmental contexts that are not part of the diurnal regime, a trend also observed in land plants and several microalgae [4]. Considering the period of rhythmic genes, genes following a 24-h period were associated with older phylostrata than genes following a 12-h period (Fig. 2a). Moreover, 24-h periodic genes peaking at ZT2, ZT6, ZT10 and ZT14 were significantly enriched in eukaryotic age, whereas genes peaking at ZT18 and ZT22 were associated with diatom and/or species-specific ages as well as genes with unknown functions (Fig. 2a). Notably, no such association between phasing and gene age was found in glaucophyta, red or green algae [4]. Taken together, genes phasing during the day and at the start of the night tend to perform deeply conserved eukaryotic functions, while previously undescribed processes specific to diatom biology appear to be occurring at midnight and during the second half of the night.

### 3.4 Rhythmic gene expression is relatively conserved between two pennate species with different lifestyles

A comparative analysis based on orthology was carried out to explore the conservation of gene expression phasing in diatoms. We restricted our comparison to the nutrient replete dataset from the *P. tricornutum* iron availability study of Smith et al. [10], as the *T. pseudonana* diurnal studies only cover two time points per day [7, 8]. We inspected the presence of phase shifts between *S. robusta* and *P. tricornutum* in 575 rhythmic one-to-one ortholog pairs with a 24-h period and found a significant association between the phasing of orthologs (Fig. 3a). The majority (60%) of *P. tricornutum* genes phased at a comparable time of the day in both species, i.e. 1h earlier or 3h later than their *S. robusta* orthologs (Chi-square Goodness of Fit test, p < 2.2e-16, std. res. of 14.7 and 2.8 respectively, Fig. 3a, Suppl. Fig. 5). Inspecting the ortholog pairs classified into each combination of phases revealed that genes with small phase shifts are involved in metabolic processes such as ribosome biogenesis, photosynthesis, proteasomal degradation and respiration, while cell cycle regulators were generally expressed 5h later in *S. robusta* (Fig. 3b).

**Figure 3:**
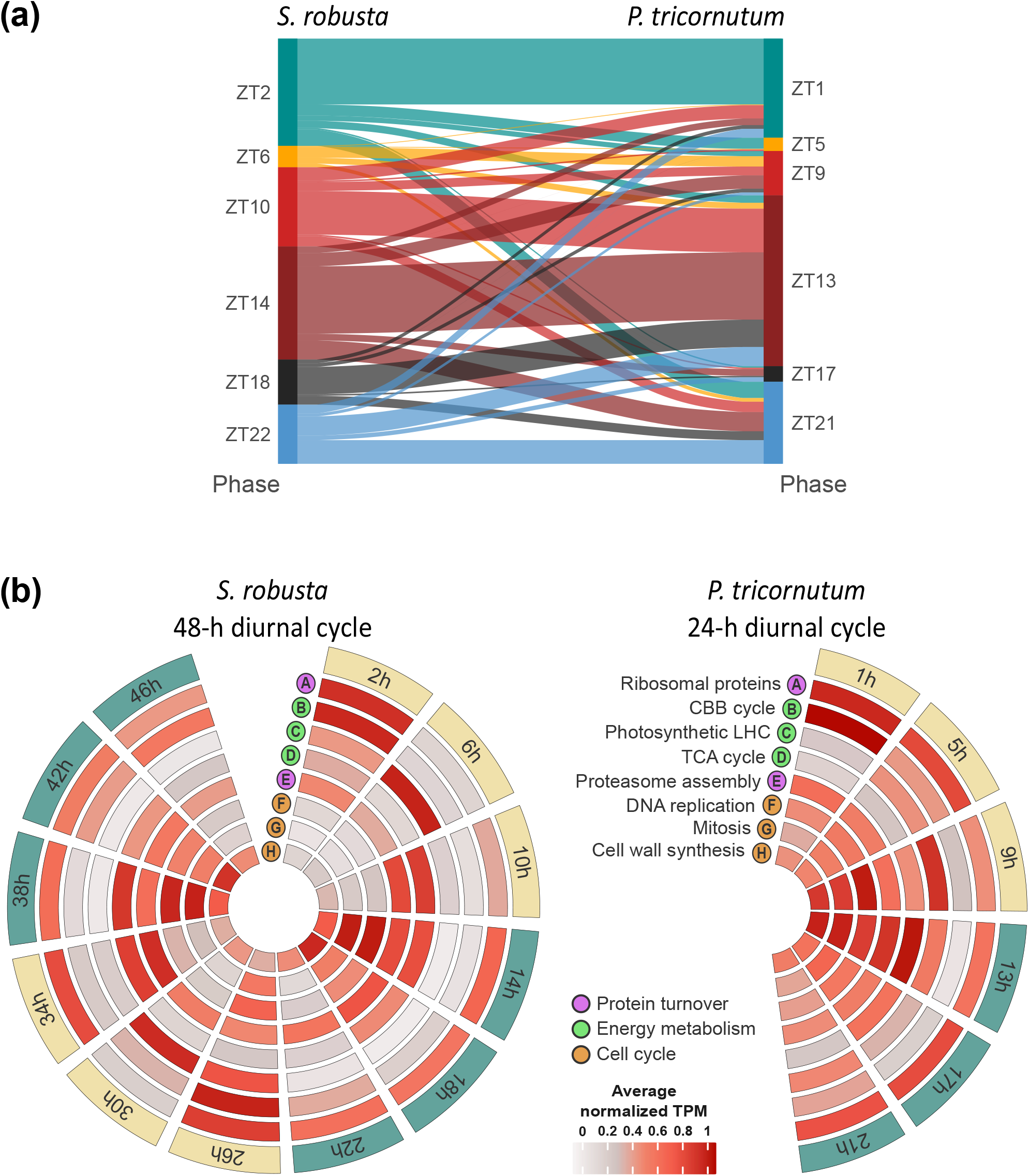
Comparison of diel expression between S. *robusta* and *P. tricornutum*. **(a)** Sankey diagram of the phase of 575 one-to-one orthologs in *S. robusta* versus *P. tricornutum*. The number of ortholog pairs within a given combination of phases is shown by the width of the line connecting the phases. The phase of the *S. robusta* ortholog (in Zeitgeber Time, h since illumination) is coded by their colour. **(b)** Circular heatmap plots displaying in every row the average normalized expression (transcripts per million, TPM) of diurnally expressed genes belonging to selected processes in *S. robusta* and *P. tricornutum*, in a color gradient ranging from grey (0) to dark red (1). In every column the time point (h) in the diurnal cycle is shown (green = dark, yellow = light). The *P. tricornutum* time point 25h represents the equivalent 1h in this figure. CBB refers to Calvin-Benson-Bassham, LHC to light harvesting complex and TCA to tricarboxylic acid cycle.

The similar phasing between the lowly silicified planktonic diatom *P. tricornutum* and the motile benthic *S. robusta*, despite considerable differences in lifestyle, indicates that the temporal regulation of basal metabolic pathways is relatively conserved.

### 3.5 Daytime anabolic and nocturnal catabolic programs

Comparative orthology analysis with *P. tricornutum* indicated a generally similar timing of metabolic processes between both species (Fig. 3b). To gather a more detailed overview of the respective timing of specific processes, we next analyzed the diel expression of well-known metabolic pathways involved in protein turnover, photosynthesis and respiration in the diurnal *S. robusta* protein-coding gene dataset.

Rhythmic expression of genes related to protein turnover was strongly coordinated in *S. robusta* (Fig. 4a, Suppl. Fig. 6). While tRNA-loading aminoacyl tRNA-ligases peaked early at ZT22, translation elongation and initiation factors were phasing at ZT2 (Fig. 4a). Expression of translation machinery at the end of the night or at dawn has been observed in the diel transcriptome of other diatoms, but also distantly related microalgae such as *Ostreococcus tauri* and *Nannochloropsis oceanica* [7, 9, 10, 49, 50]. Following protein synthesis, we found that both vesicle-mediated transport and proteasome-mediated proteolysis were enriched at dusk (Fig. 4a). The phasing of proteasomal proteolytic machinery at dusk might point towards cell cycle or circadian clock control by ubiquitination [56, 57], or may simply reflect cyclic protein turnover in which proteins are translated during the day and are recycled at night, as reported in other diatoms [7, 9, 10]. Interestingly, we observed an enrichment of 12-h periodic genes for the GO terms “translation” and “proteolysis” (31 and 64 genes respectively), the former phasing at ZT2 as well as ZT14, the latter at the second half of each day and night (Fig. 4a). While the set of 24-h period genes mainly consisted of proteasome-mediated proteolytic factors, 12-h periodic proteolytic genes contained putative lysosomal peptidases, pointing towards a temporal separation between the two main pathways for protein degradation in *S. robusta* [58].

**Figure 4:**
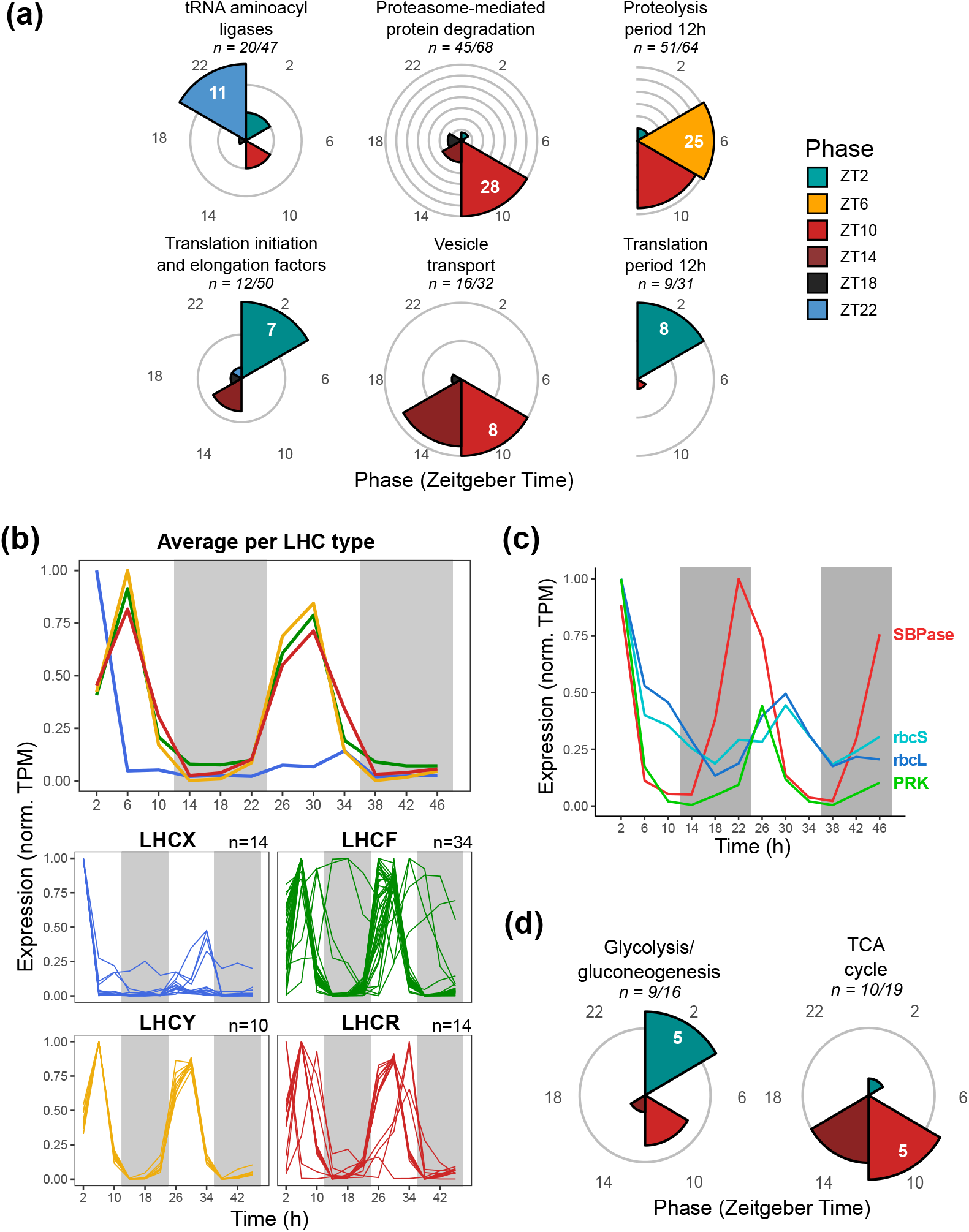
Diel expression of genes from the basal cellular metabolism. **(a)** Circular bar plots summarizing the timing of gene sets related to protein turnover. The number of genes classified into each phase cluster is indicated by the radius of each bar, with each concentric grey background circle representing an increase of four genes. The number of genes classified in the largest set is shown in white. “n” reports the number of genes with a consistent peak out of all rhythmic genes in a given category. The colour shade and the axis numbers surrounding the plot indicate the phase in Zeitgeber Time (hours since first illumination). The phase of gene sets with a 12-h (semidiurnal) period for proteolysis and translation is indicated on semicircles. **(b)** Line graph showing the expression of S. robusta light harvesting complex (LHC) genes in normalized transcripts per million (TPM) over time (in h). The upper panel shows the average expression per clade, while the lower panels show individual LHC genes. Colours indicate the four families of LHC proteins recognized in diatoms, as defined by phylogenetic analysis in Suppl. Fig. 7. The number of genes plotted is indicated by “n=“.Grey background shading represents darkness. **(c)** Line plot showing normalized expression (TPM) over time for four genes committed to the Calvin-Benson-Bassham (CBB) cycle: Rubisco (RbcS, RbcL), sedoheptulose-bisphosphatase (SBPase) and phosphoribulokinase (PRK). **(d)** Circular bar plots summarizing the timing of gene sets related to cellular respiration. For glycolysis/gluconeogenesis, only cytosolic and mitochondrion-targeted isozymes were included. TCA refers to tricarboxylic acid cycle. This panel follows the same style and colour code as panel a.

Genes involved in autotrophic metabolism were consistently phasing at dawn and midday (Fig. 4b-c). GO terms describing biosynthesis of chlorophyll and carotenoid pigments were enriched in genes peaking at ZT2, indicating that pigment biosynthesis is triggered by light onset for photosynthesis. In addition, all 73 expressed *S. robusta* light harvesting complex antenna proteins (LHC) showed significant diel rhythms. Phylogenetic analysis classified LHC genes into four distinct LHC clades: the light-harvesting fucoxanthin chl a/c LHCF, LHCR and LHCY and the photoprotective LHCX (Suppl. Fig. 7) [41, 59]. In contrast to pigment synthesis phasing at dawn, we observed a sharp peak in expression of LHCF, LHCR and LHCY genes at ZT6, likely in preparation for chloroplast division [10, 38] (Fig. 4b, Suppl. Fig. 8). An expansion of photoprotective LHCX is thought to allow benthic diatoms to cope with exposure to highly variable irradiances [41]. Here, members of the LHCX family showed an intense joint expression peak at ZT2 the first day but lacked a second-day peak in expression, even though experimental conditions were equal to the first day. Photosynthetic electron transport chain components from both photosystem I and II were enriched in genes peaking during the day, the former more specifically at ZT2. In addition, Calvin-Benson-Bassham (CBB) cycle genes [49] such as the CO_2_ capturing enzyme rubisco (rbcS and rbcL), sedoheptulose-bisphosphatase (SB-Pase) and phosphoribulokinase (PRK) phased at dawn (Fig. 4c). Genes encoding key enzymes in the biosynthesis of the storage polysaccharide chrysolaminarin were phasing at dawn as well (Suppl. Fig. 9). This is in line with a daytime accumulation and nocturnal depletion of chrysolaminarin in the diatom *P. tricornutum* [60]. The expression pattern of chrysolaminarin synthesis genes closely resembles phasing of storage compound producing pathways in other unicellular photosynthetic species: *N. oceanica* (lipids, laminarin), *O. tauri* (starch) and cyanobacteria (glycogen) [49, 50, 61].

In antiphase with photosynthetic and anabolic genes, tricarboxylic acid (TCA) cycle enzymes showed a coherent expression pattern peaking at dusk (ZT10 and ZT14). Thus, in addition to the absence of photosynthetic oxygen production at night, the dominance of cellular respiration at this time further contributes to the nocturnal depletion of oxygen in benthic ecosystems [62, 24]. Peak expression of glycolytic isozymes was incoherent and dependent on which organelle the gene product is targeted to (Fig. 4d, Suppl. Fig. 10). Apart from isozymes localized in the cytosol and chloroplast, several enzymes from the pay-off phase of the glycolytic pathway were targeted to the mitochondrion, a unique feature of stramenopile metabolism that is thought to allow simultaneous flux of incompatible reactions [63, 64, 65]. Cytosolic isozymes were either phasing at dawn or dusk, presumably supporting gluconeogenesis during the day and supplying pyruvate for respiration at night [10]. On the other hand, chloroplast and mitochondrial-targeted glycolytic isozymes were co-expressed at ZT2 the first day (Suppl. Fig. 10), substantiating the existence of a temporal separation of glycolytic pathways and a close coupling between mitochondrial glycolysis and the CBB cycle in diatoms [8, 10].

The phasing of autotrophic and heterotrophic pathways is remarkably similar between *S. robusta* and available diurnal transcriptomes from *N. oceanica* (stramenopile), *Cyanophora paradoxa* (glaucophyte), *Chlamydomonas reinhardtii* and *O. tauri* (green algae) [3, 4, 49, 50]. In these distantly related microalgae, CBB cycle genes are also phasing at dawn or during the first half of the day, followed by light harvesting genes peaking at midday and TCA cycle enzymes at nightfall. Even the cyanobacteria *Prochlorococcus* and *Synechocystis* follow the same diurnal sequence [61, 66, 67]. Carbohydrate pathways are among the most rhythmic gene sets in the transcriptome of unicellular algae, while they are largely arrhythmic in multicellular plants [4].

### 3.6 Marker genes and physiological measurements reveal rapid nocturnal progression through the cell cycle

The *S. robusta* cell cycle was highly synchronized to the diel cycle, supported by 86% (176/204) of cell cycle marker genes being significantly rhythmic and the oscillation of physiological measurements of cell division (Fig. 5a). Coordinated cell cycle progression during the diel cycle has been observed in all major groups of microalgae, suggesting an evolutionary advantage such as the temporal separation of growth and cell division [4, 49, 68, 69]. The expression of the *S. robusta* homolog of the *P. tricornutum* G1 re-entry gene dsCyc2 at ZT2 implies that a majority of cells start the day in the G1 phase while the G1-associated cyclin-dependent kinase CDKA1 phased at ZT10 [70, 71]. G1/S transition and S phase marker genes were phasing at ZT14 with an additional peak the second day at ZT2, the latter indicating a secondary round of cell cycling, a subpopulation dividing during the day or organelle DNA replication (Fig. 5a). Although care was taken to avoid nocturnal light contamination, we cannot exclude that the additional S phase peak is caused by the presence of very low background light the first night. Flow cytometry revealed that each peak in S phase expression is followed by an increase in cells with a duplicated genome 4h later (Fig. 5a “G2/M phase cells”). Following genome replication, chromosome assembly markers and the mitotic cyclin-dependent kinase CDKA2 [72] were maximally expressed at ZT14, while markers for metaphase-to-anaphase transition such as the anaphase promoting complex APC/C activator cdc20 [73] phased at ZT18 (Fig. 5a). Cell division was mainly restricted to the night, with the proportion of doublets (dividing cells) peaking at ZT18 and ZT22 and cell density almost doubling overnight (80% increase, Suppl. Fig. 11). Remarkably, while S phase markers and flow cytometry indicated a secondary round of cycling restricted to the second day, such an extra peak was not supported by mitotic and cytokinetic marker genes or an increase in the cell number (Fig. 5a, Suppl. Fig. 11).

**Figure 5:**
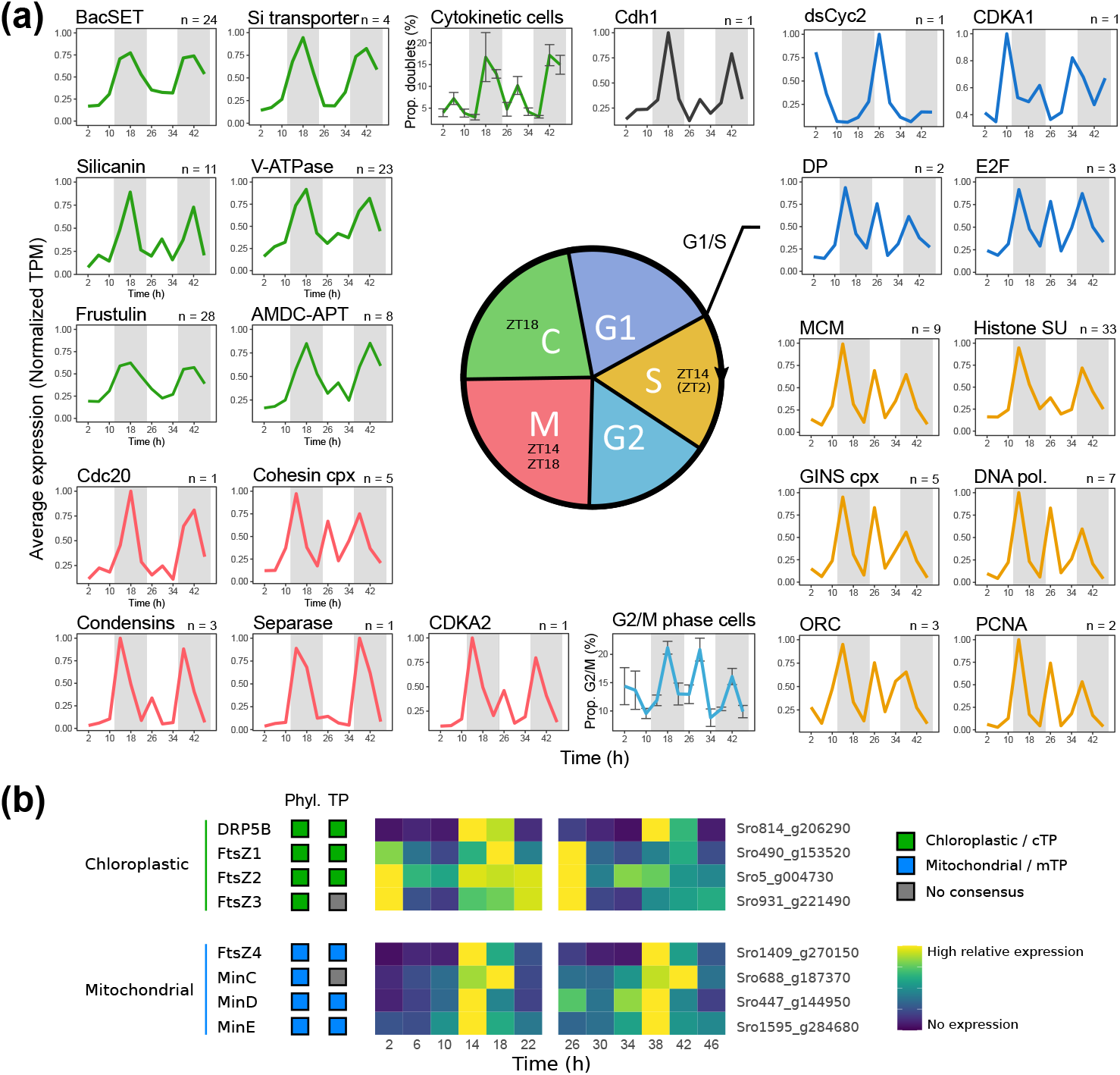
Diel transcriptional dynamics of marker genes for cell cycle and organelle division. **(a)** Line plots showing the average normalized expression (transcripts per million, TPM) over time (h) of significantly rhythmic marker genes for cell cycle progression. The number of rhythmic marker genes averaged in each plot is shown in the top right corner (n). The colour of the line refers to the cell cycle stage as indicated in the central cartoon, where also the consensus phase of marker gene expression for different stages is indicated in Zeitgeber Time (hours since first illumination). The timing of an additional peak uniquely occurring during the second day is shown between brackets. Cell cycle stages in the cartoon are not necessarily proportional to the length of these stages in the experiment. Line plots depicting the average percentage of G2/M phase cells (“G2/M phase cells”) and microscopically visible cytokinetic cells (“Cytokinetic cells”) are included, with error bars representing the standard error. DP = Dimerization Partner, MCM = Minichromosome maintenance complex, SU = Subunit, cpx = complex, pol. = polymerase, ORC = Origin recognition complex, PCNA = Proliferating cell nuclear antigen, AMDC-APT = S-adenosylmethionine decarboxylase - aminopropyltransferase family, BacSETs = Bacillariophyceae-specific SET domain protein methyltransferases. **(b)** Classification and diel expression of organelle division genes. Coloured boxes on the left show the predicted target organelle (chloroplast = green, mitochondrion = blue) based on gene phylogeny (“Phyl.”) and the presence of an N-terminal transit peptide (“TP”). TPM normalized to a maximum of 1 over each day is shown as a heatmap for each of these genes, following a color gradient from blue (no expression) to yellow (maximal expression). cTP = chloroplastic transit peptide, mTP = mitochondrial transit peptide.

A preference for genome replication taking place around dusk is observed in most lineages of microalgae and is thought to avoid DNA damage by UV or photosynthetic reactive oxygen species (ROS) during this sensitive phase of the cell cycle [74, 75]. This is reflected in the expression of S phase genes in the afternoon and at dusk in transcriptomes of *N. oceanica*, *O. tauri*, *P. tricornutum*, and *C. reinhardtii* [3, 9, 10, 49, 50, 68]. Subsequently, mitosis and cell division in these species take place at dusk or during the first half of the night. In *S. robusta*, the expression of cell cycle markers appears to be slightly delayed compared these other microalgae: genes encoding S phase machinery peaked at ZT14 (compared to ZT9 in *P. tricornutum*) and cytokinesis predominantly occurred at ZT18 and ZT22 (Fig. 3b “F”, Fig. 5a). Given the relatively low irradiance used in our experiment, the delay of S phase onset might be explained by the existence of a G1 phase commitment point which allows progression through the cell cycle only when sufficient photosynthetic energy has been captured. A dependence of S phase timing on light intensity in the pre-commitment period has been found in green algae but such a commitment point has not yet been demonstrated in diatoms [76, 77]. In addition, cell cycle progression in natural conditions might be restricted until cells have migrated at dusk, as was observed for several other species of motile benthic diatoms [25, 51].

### 3.7 Diurnal cycles in chloroplast and mitochondrion division machinery

A combination of phylogenetics and transit peptide prediction revealed that genes encoding bacterial division-like proteins were targeted to the chloroplast and the mitochondrion (Fig. 5b, Suppl. Fig. 12-13), pointing to the retention of an ancestral system for mitochondrial division in *S. robusta*, as was observed in various proists, including diatoms but with the notable exception of *P. tricornutum* [78, 79, 80]. While the eukaryotic chloroplast division gene DRP5B (Suppl. Fig. 14) was phasing at ZT14 in accordance with autonomous chloroplast division being coordinated to the S-G2 phases of the cell cycle [38], all three bacterial-derived chloroplastic FstZ genes showed a maximum expression at ZT2 with additional expression at night (Fig. 5b). The discrepancy between the timing of FtsZ expression and chloroplast division in *S. robusta* is highly unusual, given that the FtsZ peak of expression is coordinated with chloroplast division in other microalgae such as *O. tauri* and *Cyanidioschyzon merolae* [50, 81]. Moreover, FtsZ and DRP5B peaked simultaneously in the afternoon in *N. oceanica*, and chloroplast division was coordinated with nuclear division in the related species *Nannochloropsis oculata* [82, 49]. It is worth noting that the stramenopile chloroplast is surrounded by four membranes, of which the outermost is continuous with the nuclear envelope [83]. Fission of the outer chloroplast membrane of the pennate diatom *P. tricornutum* only occurs after the three inner membranes have bisected the chloroplast [83]. This suggests an independent (FtsZ-free) division mechanism for the outermost membrane, and implies a coordination with nuclear division like in *N. oculata* [82].

All mitochondrial FtsZ and Min-system division genes were significantly rhythmic and showed a consistent daily burst at ZT14 and ZT18 (Fig. 5b), suggesting that mitochondria either divide in coordination with maximal respiration at night, or with the cell cycle as was observed in the red alga *C. merolae* [81] and suggested in *P. tricornutum* [84].

### 3.8 Tight coordination of cell division over the diurnal cycle uncovers potential uncharacterized cell wall biosynthetic genes

The peak of cytokinesis at ZT18/ZT22 coincided with the expression of rhythmic genes involved in cell wall formation, which takes placed in a specialized intracellular compartment termed the silica deposition vesicle (SDV): 23 SDV-acidifying V-type ATPases, 11 SDV transmembrane silicanin-like genes, 24 diatom-specific SET domain protein methyltransferases (BacSETs), eight S-adenosylmethionine decarboxylase-aminopropyltransferase (AMDC-APT) genes, four silicic acid transporters and 28 genes from the frustulin family were phasing predominantly at ZT18 (Fig. 5a). Notably, an 11-transmembrane *S. robusta* homolog of the plant Silicon efflux transporter Lsi2 was also co-expressed at ZT18. This gene was downregulated during nutrient depletion and pheromone induced cell cycle arrest, supporting a role as an additional type of silicic acid transporter in diatoms as was hypothesized in *T. pseudonana* (Suppl. Fig. 15) [85]. The majority (82/113) of the cell wall genes was of diatom-specific age in our phylostratigraphic analysis, indicating that silica cell wall biosynthetic machinery might have evolved de novo in a common ancestor of the diatoms and the pico-eukaryotic sister group Bolidophyceae [86]. The over-representation of uncharacterized, diatom-specific genes phasing simultaneously with known cell wall formation factors suggests that hitherto unknown cell wall biosynthesis gene families are active at ZT18 (Fig. 2a). *T. pseudonana*, the best established model for cell wall biogenesis, displayed a lower expression of cell wall biosynthesis genes under silicon depletion conditions [85, 87]. Inspection of genes downregulated after silicon depletion in *S. robusta* yielded a significant association with genes phasing at ZT18 (q < 0.05). Among the top six diatom-specific families in this dataset, we observed the known cell wall associated frustulins and silicanins as well as four new candidate cell wall formation families that we termed Siren1-4 (“**Si**lica **re**sponsive **n**ight”, Fig. 6). The presence of a predicted signal peptide in Siren1 and Siren3 proteins supports a possible role in cell wall formation, as a signal peptide is required for SDV protein targeting (Suppl. Fig. 16) [88, 85]. Although Siren2 and Siren4 do not possess a characteristic signal peptide and therefore may rather be involved in other processes such as late-stage mitosis, they may also participate in cytosolic processes that are suggested to influence silica morphogenesis [89, 90]. Siren1 and Siren4 genes do not contain known protein domains, but Siren2 and Siren3 were annotated with ankyrin repeats and an alpha/beta hydrolase fold domain, respectively. During cell wall synthesis, the protein-interacting ankyrin repeats in the Siren2 family may regulate cytoskeletal organization or cell cycle progression, processes these motifs are often involved in [91]. Over the years, numerous proteins associated with diatom cell wall biosynthesis (e.g. silaffins, cingulins and silacidins) have been proposed, but only a handful have a broad distribution across diatoms, most notably the silicanins, frustulins, p150 proteins and BacSETs [88]. We expand this list with the putative Siren3 and Siren4 cell wall families which are shared among pennate and centric diatoms (Fig. 6). Homologs of Siren1 and Siren2, on the other hand, appeared to be uniquely present in pennate diatoms, which is exciting as no pennate-specific cell wall associated proteins have been described to date (Fig. 6) [88].

**Figure 6:**
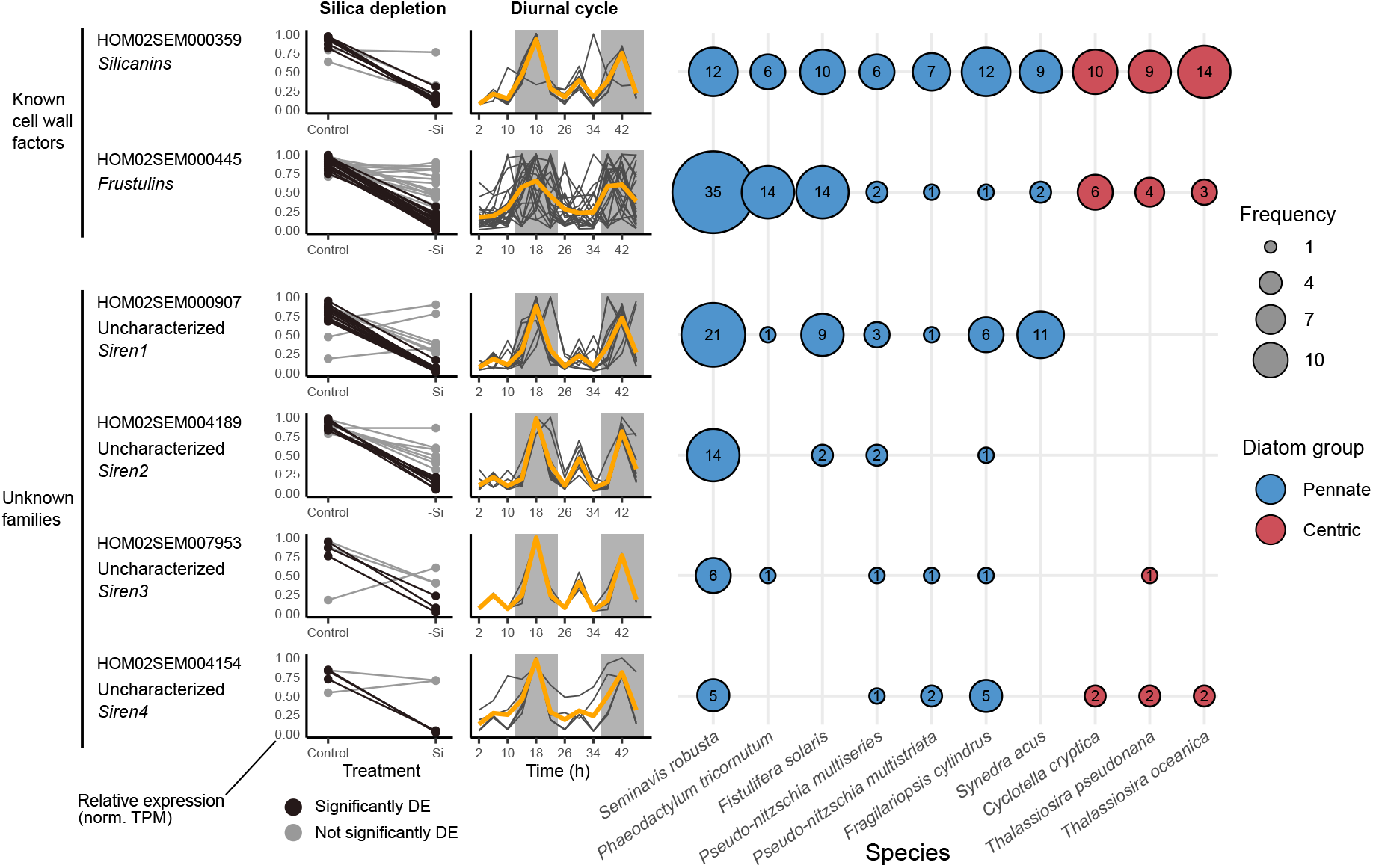
Identification of putative new diatom cell wall biosynthesis families using silica depletion and diurnal cycle RNA-seq datasets. The PLAZA Diatoms 1.0 family ID and description of six gene families are given on the left hand side, with unknown families being termed “Siren” (Silica responsive night). The relative expression of members of each gene family during silica depletion and in the two-day expression dataset is shown as transcripts per million (TPM) normalized for each gene to a maximum of 1 over the experiment. Solid lines connect observations from the same gene over conditions or time. Genes significantly differentially expressed (“DE”) in response to silica depletion are plotted with black lines, while an orange line shows the average relative expression of a gene family over the diurnal cycle experiment. Diel expression was only plotted for significantly rhythmic genes with a period of 24h. On the right, the presence of members of the gene family in the genome of 10 diatom species is shown, with circle radii and numbers inside circles showing the family size in each species. Gene families in pennate diatoms are shown with a blue filled circle, while centric diatoms are indicated by a red shade.

### 3.9 Linking the expansion of putative circadian genes to benthic diatom biology

The Per/Arnt/Sim (PAS) signalling domain is a hallmark of many proteins that constitute the central circadian oscillator in animals, often alongside a basic helix-loop-helix (bHLH) DNA binding domain [92]. The bHLH-PAS transcription factors (TFs) have recently been identified as the first known components of the stramenopile circadian clock, with the discovery of the timekeeping regulator bHLH1a-PAS (RITMO1) in the diatom *P. tricornutum* [11]. In *S. robusta*, we identified 13 bHLH TFs, of which 10 contained a PAS domain, which is in stark contrast to *P. tricornutum*, encoding only three bHLH-PAS [11]. Phylogenetic analysis revealed that each *P. tricornutum* bHLH TF corresponded to one *S. robusta* ortholog, except for bHLH2-PAS which expanded into seven gene copies (Fig. 7). RITMO1 and bHLH3 orthologs showed conserved rhythmic expression, while bHLH1b-PAS displayed antiphase expression between both species (Fig. 7).

The multi-copy *S. robusta* bHLH2-PAS clade of which 6/7 were rhythmic, exhibited expression divergence, with rhythmic genes peaking either at dawn or broadly during the day (Fig. 7). The expansion and expression divergence of bHLH-PAS TFs may suggest that the inherently variable and highly rhythmic benthic environment drove the evolution of extensive time-keeping mechanisms. Additionally, two circadian-regulated *P. tricornutum* bZIP-PAS TFs (bZIP5-PAS and bZIP7-PAS) showed a conserved diel expression in *S. robusta*, phasing at night and at dusk respectively (Suppl. Fig. 17) [11]. Together, the phasing of *S. robusta* bHLH-PAS and bZIP-PAS TFs covered the complete diurnal cycle, suggesting that a sequential expression of these TFs drives the circadian clock of this species (Suppl. Fig. 17).

Two other types of presumed diatom circadian genes showed significant oscillations [93, 94] (Suppl. Fig. 18). We observed a peak at night for three casein kinase 1 (CK1) homologs while the blue-light sensing cryptochrome/photolyase family 1 (CPF1) gene was peaking at dawn, analogous to *P. tricornutum* and *N. oceanica* [49]. Genes encoding G-protein-coupled receptor (GPCR) rhodopsin-like proteins, which are prominently expanded in the *S. robusta* genome and conserved in several other benthic diatoms, were predominantly expressed at dusk and at night (Suppl. Fig. 19) [33]. Given the sensitivity of animal rhodopsins to extremely low levels of light, they may be involved in photoreception inside sediments or phototrophic biofilms which exhibit strong light attenuation on a millimeter scale [95, 96, 97]. Alternatively and in accordance with their nocturnal expression, GPCR rhodopsin-like proteins may be used to perceive twilight or moonlight at night, a feature observed for rhodopsins in the compound eyes of insects [98, 99].

### 3.10 Rhythmic genes with a 12-h period encode putative exopolymer production enzymes

Next to diurnally regulated genes, we observed that 7.5% of rhythmic genes displayed a semidiurnal rhythm, i.e. had a period of 12h (Fig. 1c). Apart from translation factors and proteases/peptidases, we observed that a range of glycosyltransferases, sulfotransferases, sugar phosphate transporters and fasciclin-like adhesive proteins are enriched in 12-h periods and phase at ZT10/ZT22. In diatoms, such genes have been implicated in the production of EPS, which are typically produced during migration and adhesion [100, 101, 102]. The semidiurnal expression of these genes might be linked to twice-daily vertical migration in subtidal and intertidal diatom communities, where diatoms burrow into the sediment at dusk and emerge at dawn [24, 25, 103]. Alternatively, semidiurnal rhythmicity may be an adaptation to the occurrence of tides in coastal benthic ecosystems, as there is typically a period of 12.4h from one high tide to the next one, and tides were observed to induce cyclic migration patterns in intertidal diatoms [24, 104, 105]. In several coastal animals, gene expression with a 12-h period has already been attributed to tidality [106, 107]. Nonetheless, our laboratory experiment did not allow for vertical migration and did not simulate tidality, making it hard to attribute this semidiurnal phenomenon to any of these scenarios. Finally, a hybrid-sensor PAS (Sro442_g143910) and a bZIP transcription factor (Sro199_g084500, bZIP1) showed significant 12-h oscillations phasing at ZT10/ZT22, both being credible candidate components of a potential circatidal clock as has been described in marine animals [93, 106, 108, 109].

### 3.11 Concluding remarks

Comprehensive profiling of the *S. robusta* transcriptome revealed that oscillating proteincoding genes are organized into highly coordinated transcriptional programs controlling cell cycle progression, protein turnover and carbohydrate metabolism. Moreover, the identification of rhythmic lincRNAs indicates that the non-coding transcriptome is not solely important in the response to environmental stressors in diatoms [47, 48], but also plays a role in the diurnal cycle. The pronounced differences in expression levels of these processes over the course of the day should be considered when interpreting data from metatranscriptome initiatives such as TARA Oceans where samples are collected at a particular time of the day and at varying latitude with inherently different day lengths, as diel rhythms will likely have a profound impact on gene expression in each sample [110, 111].

**Figure 7:**
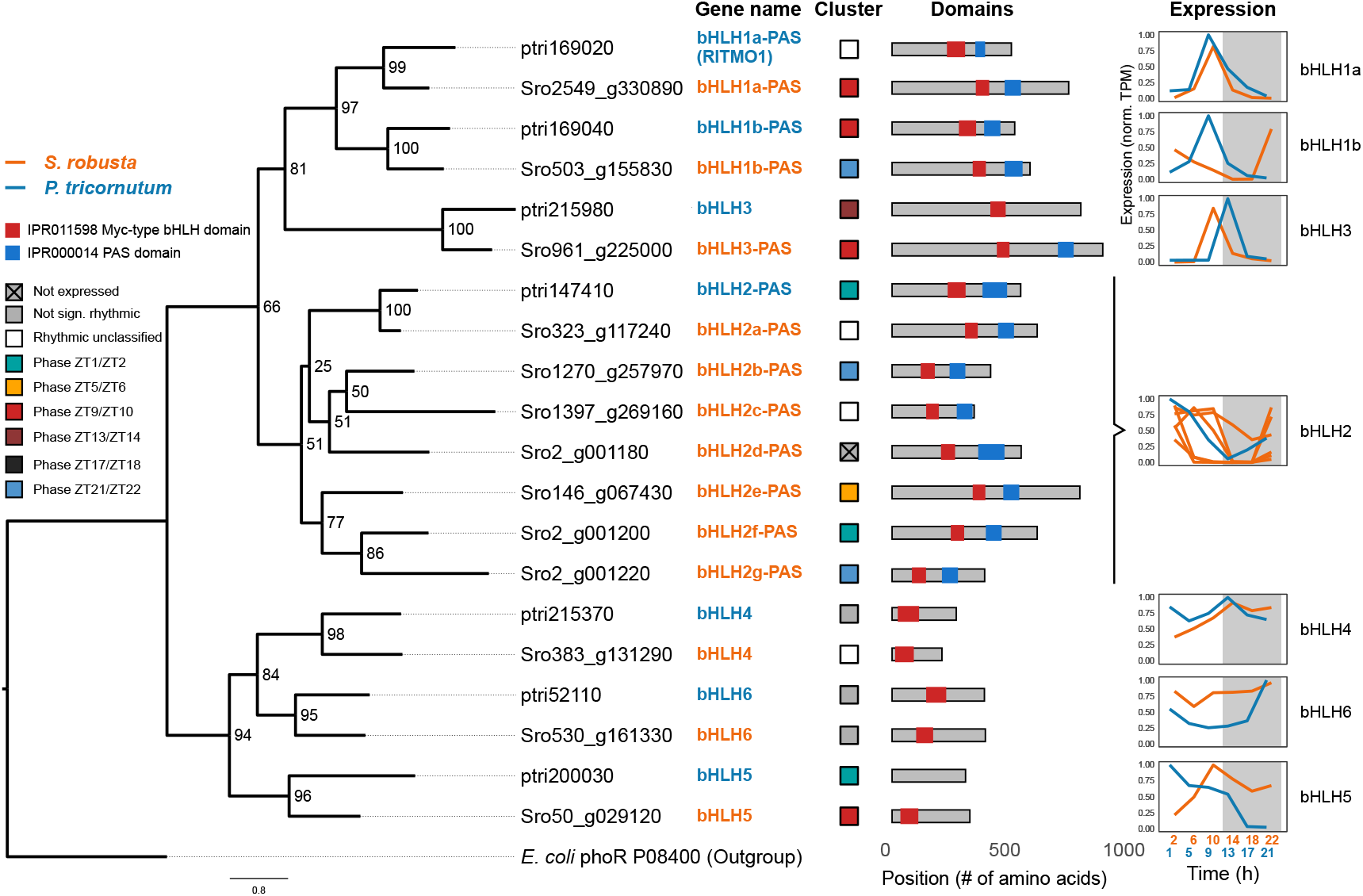
Phylogeny and comparative analysis of the diel expression of basic helix-loop-helix (bHLH) transcription factors in the pennate diatoms *S. robusta* and *P. tricornutum*. The left side shows the maximum likelihood phylogenetic tree of bHLH transcription factors, with bootstrap support values shown at the branch points. The *E. coli* PAS-domain containing protein PhoR was used as an outgroup for rooting. For each gene in the tree, coloured boxes indicate the classification of diurnal oscillations (“cluster”): not expressed, not significantly rhythmic, rhythmic with unclassified phase and a specific phase cluster (in Zeitgeber Time, hours since illumination). The position of bHLH (red) and PAS (blue) InterPro domains on each gene (grey) is visualized under “Domains”. When multiple identical overlapping domains were predicted, the longest domain was selected for visualization. Average normalized diel expression (transcripts per million, TPM) over time (in hours) is shown by line plots for each subclade at the right side (*S. robusta* = orange, *P. tricornutum* = blue). To allow comparison with the 1-day *P. tricornutum* time series, *S. robusta* relative expression was averaged over both days and shown over the span of one day. The *P. tricornutum* time point 25h represents the equivalent 1h. Grey shaded areas represent darkness treatment.

The diurnal regulation of carbohydrate metabolism appears to be conserved among microalgae. Not only was autotrophic and heterotrophic transcription temporally segregated as the result of photosynthetic energy input being restricted to the day, we also observed anticipatory expression of anabolic and photosynthetic genes before sunrise, which suggests regulation by the circadian clock [7, 50, 61].

The strict diurnal orchestration of gene expression patterns and the behaviour of *S. robusta* cells revealed new potential cell wall biosynthetic families, a spatial and temporal segregation of the glycolytic pathway, and indications of synchronous mitochondrial division. In addition, the expansion of circadian transcription factors and photoprotective LHCX, and the 12-h periodicity of putative EPS-producing enzymes all suggest that raphid pennate diatoms evolved multiple adaptations to the complex and variable benthic environment. Additional transcriptomic profiling of benthic diatoms would benefit from a set-up allowing for vertical migration while simulating tidal regimes. Similarly, the global transcriptomic response to different photoperiods has not been assessed in diatoms despite it being a highly relevant topic, as daylength at high latitudes varies greatly throughout the year due to seasonality, and diel expression of circadian regulators in diatoms is sensitive to these changes in periodicity [11].

## 4 Methods

### 4.1 Experimental conditions of the diel experiment

*Seminavis robusta* strain 85A (DCG 0105), obtained from the Belgian Coordinated Collection of Microorganisms (BCCM/DCG, http://bccm.belspo.be/about-us/bccm-dcg), was grown in artificial sea water (ASW) prepared using 34.5 g/L Tropic Marin BIO-ACTIF sea salt (Tropic Marin, Wartenberg, Germany) and 0.08 g/L sodium bicarbonate. After autoclaving, ASW was supplemented with Guillard’s F/2 Marine Water enrichment solution (Algoid technologies) and an antibiotic mix (400 μg/ml penicillin, 400 μg/ml ampicillin, 100 μg/ml streptomycin and 50 μg/ml gentamicin) [112]. Cultures were grown in 150 mL of medium in Cellstar Standard Cell Culture Flasks (Sigma-Aldrich) with a surface area of 175 cm^2^, which were placed in a climate-controlled plant growth chamber (Weiss technik) at a temperature of 21°C and a light intensity of 30 μE m^−2^ s^−1^. Cultures were first acclimated to a 12h-light/12h-dark photoperiod for 5 days, after which they were sampled for transcriptomic analysis at 12 time points spanning two diurnal cycles (day, night, day, night). The first culture flask was harvested the first day after 2h of illumination (ZT2). Subsequently, samples were harvested every 4h, resulting in 6 samples for the first cycle (2h, 6h, 10h, 14h, 18h, 22h) and 6 samples the next (26h, 30h, 34h, 38h, 42h, 46h). At each time point, a complete culture flask was harvested by scraping cells from the bottom surface using a cell scraper and homogenizing by gentle shaking. Cells were collected by filtration with a pore size of 3μm and flash frozen in liquid nitrogen to be stored at −80°C awaiting RNA extraction. For samples collected at night, this procedure was performed in the dark to avoid responses to light. The experiment was repeated three times on separate occasions so that each time point was sequenced in biological triplicate (36 samples).

### 4.2 Quantification of cell cycle progression

To assess S phase progression during the experiment, a subsample of RNA-seq cultures were analysed using flow cytometry (Suppl. Fig. 20): 5 mL of suspended culture from each 150 mL culture flask harvested for RNA-seq was held apart for flow cytometry analysis. Next, this 5 mL of culture was centrifuged for 5 min at 3,000 RPM. The pellet was fixated by adding 10 mL of ice cold 75% ethanol, after which samples were kept in the dark at 4 °C. Preparation of samples, staining with SYBR green and flow cytometry analysis on a Bio-Rad S3e cell sorter was performed as detailed by Bilcke et al. (2021) [32]. In addition, before harvesting for RNA-seq, microscopic pictures were taken of the culture flasks with a Canon Powershot G11 camera mounted on a Primovert inverted microscope at 20x magnification. At night, pictures were taken from a separate culture that was not afterwards used for RNA sequencing in order to not disturb the diurnal cycle. These pictures were used to determine the cell density and proportion of cytokinetic cells at each time point. Manual cell counting was performed using ImageJ software. We defined cytokinetic cells as cells showing a visible cleavage furrow or developing frustule subdividing the daughter cells.

### 4.3 RNA preparation and sequencing

Total RNA was prepared using the RNeasy Plant Mini Kit (Qiagen). Culture was scraped from the filter into 1 mL of RLT+ME buffer mixed with 1.0 mm diameter SiC beads (BioSpec products) and were lysed on a beating mill (Retsch) for 1 min at 20 Hz. The lysate was transferred to a spin column and the rest of the protocol was performed according to manufacturer’s instructions. RNA concentration was determined on the Nanodrop ND-1000 spectrophotometer (Thermo Scientific) and RNA integrity was assessed using a BioAnalyzer 2100 (Agilent). Library preparation was performed using the Illumina TruSeq Stranded mRNA Sample Prep Kit (Illumina), and included mRNA enrichment using poly-T oligo-attached magnetic beads. Paired-end 2×76 bp sequencing of 36 samples was performed on the Illumina HiSeq 4000 platform at the VIB Nucleomics core (www.nucleomics.be).

### 4.4 Long intergenic non-coding RNA (lincRNA) gene annotation and conservation analyses

LincRNAs were annotated in the *S. robusta* genome leveraging several RNA-seq datasets. The quality of adaptor-trimmed reads belonging to 161 samples from the previously published *S. robusta* gene expression atlas (Suppl. Table 2) [33] and the 36 new diel cycle samples was assessed using FastQC v0.11.4 [113]. All reads were mapped to the *S. robusta* genome using STAR v2.5.2 (–sjdbGTFfeatureExon, –alignIntronMax 6000) [114, 33] followed by the removal of chimeric alignments and retention of uniquely mapping reads using samtools v1.3 (-F 4 -q 1) [115]. Next, these alignments were split per scaffold and genome-guided Trinity De novo Transcriptome v2.6.6 was executed (–genome_guided_bam, –genome_guided_max_intron 6000) [116], generating 221,735 transcripts. Selecting the major isoform of each transcript led to an initial set of 143,473 distinct transcript models which were mapped back to the *S. robusta* genome using gmap v2017-11-15 (-min-identity 0.9) [117]. We focused our analysis on lincRNAs, whose expression quantification is the most reliable since they do not overlap with protein-coding gene regions. Therefore, transcripts overlapping with protein-coding genes were removed using bedtools v2.2.28 (-intersect) [118], retaining 33,849 transcripts. Afterwards, the FEELnc v2018-07-12 pipeline was used to annotate lincRNA candidates [119]. In detail, we executed the FEELnc filter (–monoex=1) removing transcripts with a length of <200nt and bi-exonic transcripts having one exon with a length of <25nt. Subsequently, we executed FEELnc codpot with –mode=intergenic and –mode=shuffle parameters. Transcripts with coding potential in any of these two runs were discarded (30,968 transcripts were kept). Transcripts suspected to be fragments of 5’ or 3’ UTRs, i.e. ending close to a protein-coding gene (distance <317 nt, average UTR length of protein-coding genes) were removed, retaining 18,262 transcripts. Additionally, potential remaining transcript redundancy was removed using the cuffmerge tool from cufflinks v2.2.1 (default parameters) [120]. Finally, only transcripts without similarity to protein-coding genes were retained by comparing transcripts to known proteins in the Uniprot database using blastx v2.6.0 [121, 122]. LincRNA transcripts were discarded if their E-value was below a threshold value of 1.23e-05 that was experimentally determined as the minimal blastx E-value of ten randomly shuffled sequences per candidate transcript, obtaining a set of 14,782 candidate lincRNAs (13,470 mono-exonic and 1,312 multi-exonic). Finally, a subset of 7,117 high-confidence lincRNAs which show robust expression support (average TPM > 2 in at least one condition) in either the *S. robusta* gene expression atlas or the new diurnal expression dataset was selected for further consideration.

To assess conservation of the predicted lincRNAs in other diatoms, a blastn v2.6.0 [121] search was executed against the diatom MMETSP transcriptomes [123] and nine diatom genomes stored in PLAZA Diatoms 1.0 (https://bioinformatics.psb.ugent.be/plaza/versions/plaza_diatoms_01/). The same blastn search was executed on 10-fold shuffled sequences generated from candidate lincRNAs to determine a cut-off for significant hits (E-value < 3.8e-04).

### 4.5 Quantification of diel cycle gene expression and detection of rhythmic genes

Adaptor-trimmed reads from the 36 diel cycle samples were mapped to the longest isoform for each protein-coding gene (36,254) of the *S. robusta* genome merged with the set of candidate predicted lincRNAs (14,782) using Salmon v1.1, using following parameters for indexing: –keepDuplicates -k 31, and quantification: –seqBias –gcBias -l A [124]. To assess reproducibility between diel cycle replicates and corresponding time points over subsequent days, count estimates were imported using tximport [125] and an unsupervised multidimensional scaling plot was created as implemented in the EdgeR package for R [126]. Distances in this plot approximate the log2 fold changes between pairs of samples, based on the expression (in counts per million) of the 500 most distinctive genes.

Genes of which the diel expression in transcripts per million (TPM) averaged over the three replicates exceeded two in at least one time point were retained for further analysis, keeping 26,109 protein-coding and 1,680 lincRNA genes. Genes with a rhythmic diurnal expression were identified using the empirical Jonckheere-Terpstra-Kendall (empJTK) tool [127, 128]. Genelevel TPMs of three replicates for each of the 12 time points were used as input, specifying the waveform family as “cosine”, the possible periods of the waveform as 24h and 12h, and allowing asymmetries ranging from 4h to 20h in steps of 4h. The false discovery rate (FDR) for the set of rhythmic genes was controlled at 0.05, after adjusting p-values for multiple testing using the Benjamini-Hochberg method implemented in empJTK as “GammaBH”.

Comparisons with rhythmic gene expression of the pennate diatom *Phaeodactylum tricornutum* were performed based on publicly available SOLiD RNA-seq data retrieved from the iron depletion study of Smith et al. (2016) (Suppl. Table 3). Triplicate RPKM values for 6 time points (5h, 9h, 13h, 17h, 21h, 25h = 1h) concerning the “High” iron condition (400 pM Fe, nutrient replete conditions) were converted to TPM. After removal of duplicated, empty rows and lowly expressed genes with average TPM < 2 in all time points, 10,771 genes were used for rhythmicity detection with analogous empJTK settings as for *S. robusta*, but specifying possible phases as ZT1-ZT21 with a step size of 4.

### 4.6 Classification of rhythmic genes into clusters

Genes were classified with regard to rhythmicity in a hierarchical way. The first distinction concerns rhythmic versus non-significantly rhythmic genes, as determined by empJTK. At the next level, empJTK classified rhythmic genes as having a period of 24h (one cycle each day) or 12h (two cycles each day). Further, genes were classified according to their phase, i.e. the timing of maximum expression. Although the phase of the best fitting waveform is reported by empJTK, we further restricted the phase classification to genes with a stably recurring maximum each day, taking advantage of our two-day dataset. In detail, genes were annotated with a phase if the timing of the maximal average TPM for each day (24h period) or half day (12h period) agreed with the predicted phase by empJTK. Finally, rhythmic genes were classified into broad temporal categories if their predicted and actual expression maxima and minima fall within and outside predetermined time intervals respectively: day (ZT2, ZT6, ZT10), night (ZT14, ZT18, ZT22), dawn (ZT22, ZT2) and dusk (ZT10, ZT14). *P. tricornutum* rhythmic genes were classified into clusters following the same set of rules defined for *S. robusta*.

### 4.7 Functional annotation of proteincoding and lincRNA genes

Functional annotation of *S. robusta* proteincoding genes was obtained from PLAZA Diatoms 1.0 (https://bioinformatics.psb.ugent.be/plaza/versions/plaza_diatoms_01/). In particular, Gene ontology (GO), Kyoto Encyclopedia of Genes and Genomes (KEGG) and InterPro (IPR) domain annotation from InterProScan were retrieved [33]. Phylostratification of protein-coding genes was calculated based on the taxonomic scope of their predicted TribeMCL gene family in PLAZA Diatoms 1.0, after which genes were assigned to the oldest phylostratum they belong to [129, 130]. Differential gene expression (DE) labels for *S. robusta* protein-coding genes were retrieved from Osuna-Cruz et al. (2020) [33]. Significant enrichment of phylostrata/gene family/DE/GO/IPR features for a gene set were inferred using a hypergeometric test, setting the minimum number of positive features in a set to two and controlling p-values for multiple testing on a 5% FDR level using the Benjamini-Hochberg procedure.

Combining the diurnal expression dataset with the biological process GO terms assigned to *S. robusta* protein-coding genes, a co-expression network was constructed to annotate lincRNAs based on the guilt-by-association principle [131]. Pearson correlations were computed in a pairwise manner between all genes and Highest Reciprocal Ranks (HRR) were calculated [132, 54, 133]. The optimal HRR threshold that maximizes the weighted F1-score (harmonic mean of the precision and recall) was estimated at 12. Co-expression clusters were defined based on this cut-off and GO enrichment was executed per cluster by performing a hypergeometric test (q-value < 0.05).

### 4.8 Detection and rhythmicity of orthologous genes between *S. robusta* and *P. tricornutum*

Orthologous gene relationships between *S. robusta* and *P. tricornutum* were inferred using the PLAZA integrative orthology method [134]. Genes were considered orthologs when they were confirmed by the OrthoFinder gene family (ORTHO) and the Best-Hits-and-Inparalogs (BHIF) methods in both species, identifying 575 highly reliable ortholog pairs with a one-to-one relationship and a stable, 24-h periodic phase in both species. For each pair, the phase shift in *P. tricornutum* as compared to *S. robusta* was calculated. Since harvesting points were offset 1h in *P. tricornutum* relative to *S. robusta*, the phase shift can take on following values: +3h (−21h), +7h (−17h), +11h (−13h), +15h (−9h), +19h (−5h) and +23h (−1h). The relationship of the phase between orthologs was visualized using a Sankey diagram as implemented in the ggalluvial package for R [135].

### 4.9 Construction of phylogenetic gene trees

Maximum likelihood phylogenetic trees were computed using the following protocol: i) multiple sequence alignment of genes/families of interest using MAFFT v7.187 [136], ii) automatic editing of this multiple sequence alignment using trimal v1.4.1 (–gt 0.1) [137], iii) phylogenetic tree construction using IQ-TREE v1.7 (-bb 1000 -mset JTT,LG,WAG,Blosum62,VT,Dayhoff -mfreq F -mrate R) [138] and iv) visualization and midpoint rooting using FigTree v1.4.4 [139]. The simplified light harvesting complex protein (LHC) tree was visualized using MEGA X [140] (Suppl. Fig. 7).

### 4.10 Prediction of subcellular localization

The localization of nuclear-encoded proteincoding genes in the *S. robusta* cell was computationally predicted for organelle division genes, glycolytic isozymes and genes from proposed novel cell wall associated gene families [141]. Mitochondrial transit peptide (mTP) prediction was performed with TargetP 2.0 in non-plant mode (http://www.cbs.dtu.dk/services/TargetP/) [142], whereas signal peptides (SP) were predicted using SignalP 3.0 (http://www.cbs.dtu.dk/services/SignalP-3.0/) [143]. Bipartite targeting signals (BTS) for import into the stramenopile secondary chloroplast were identified using ASAFind (https://rocaplab.ocean.washington.edu/tools/asafind/) [144] and HECTAR [145]. Making use of the endosymbiotic origin of chloroplasts and mitochondria, localization of organelle fission genes was confirmed based on phylogenetic clustering with bacterial homologs in selected alphaproteobacteria and cyanobacteria species.

### 4.11 Identification of putative new cell wall associated gene families

The differential expression matrix depicting the response of *S. robusta* protein-coding genes to prolonged silica depletion was retrieved from the publicly available *S. robusta* gene expression atlas [33]. An enrichment test was carried out as described before, testing for an association between the phasing of genes (ZT2-ZT22) and their response to silica depletion (upregulated, downregulated, not DE) in the differential expression analysis from Osuna-Cruz et al. (2020) [33]. Next, the enrichment of gene families in combined clusters of phase and silica response was determined, and gene families enriched in the set of genes with phase ZT18 and downregulated in response to silica depletion were withheld as potential silica biogenesis families. Six key families were selected after retaining only families with an enrichment q-value < 0.001 in the set of ZT18:downregulated genes, that have genes in more than three diatom species and no representatives outside diatoms. The novelty of uncharacterized families returned by this analysis was verified by cross-referencing with a list of known cell wall families.

## 5 Data availability

Raw Illumina RNA-seq reads are available at the European Nucleotide Archive (ENA) under accession number PRJEB40442. The improved *S. robusta* genome annotation including lincRNA genes is available through ORCAE (http://bioinformatics.psb.ugent.be/orcae/overview/Semro). Additional functional and homology information concerning genes mentioned in this work is available through the diatom-PLAZA platform (https://bioinformatics.psb.ugent.be/plaza/versions/plaza_diatoms_01/).

## 6 Acknowledgements

We would like to thank Nils Kröger for his advice in validation of novel silica families. We are further grateful to Camilla Ferrari and Koen Van den Berge for helpful discussion and to Thomas Depuydt for his assistance with applying the guilt-by-association method.

G.B. and C.M.O-C. were supported by a UGent-BOF project (GOA 01G01715) and G.B. was additionally funded by an FWO aspirant grant (3F001916). N.P. has been supported by the Deutsche Forschungsgemeinschaft (PO 2256/1–1). P.B. received funds from an Erwin Schrödinger fellowship from the Austrian Science Fund (J3692-B22) and an FWO project (G0D6114N). This work was further supported with infrastructure funded by the research council of Ghent University (BOF/GOA 01G01715) as well as an EMBRC Belgium—FWO project (GOH3817N).

## 7 Author contributions

K.V., L.DV. and W.V. initiated and managed this project, while experiments were designed by K.V., L.DV., W.V., P.B., C.M.O-C. and G.B.. The RNA-seq experiments were executed by G.B. and C.M.O-C. Flow cytometry and RNA quality control was performed by G.B., the latter with the help of S.D.. LincRNA detection and functional annotation was carried out by C.M.O-C. with the help of M.S-S.. RNA-seq quantification, rhythmic gene detection, clustering, comparative transcriptomic analysis and subsequent downstream biological interpretation were performed jointly by C.M.O-C. and G.B.. N.P. assisted in the computational validation of proposed new cell wall families. The manuscript was written by G.B. and C.M.O-C. and reviewed by all co-authors.

## 8 Supplementary Figures

**Supplementary Figure 1:**
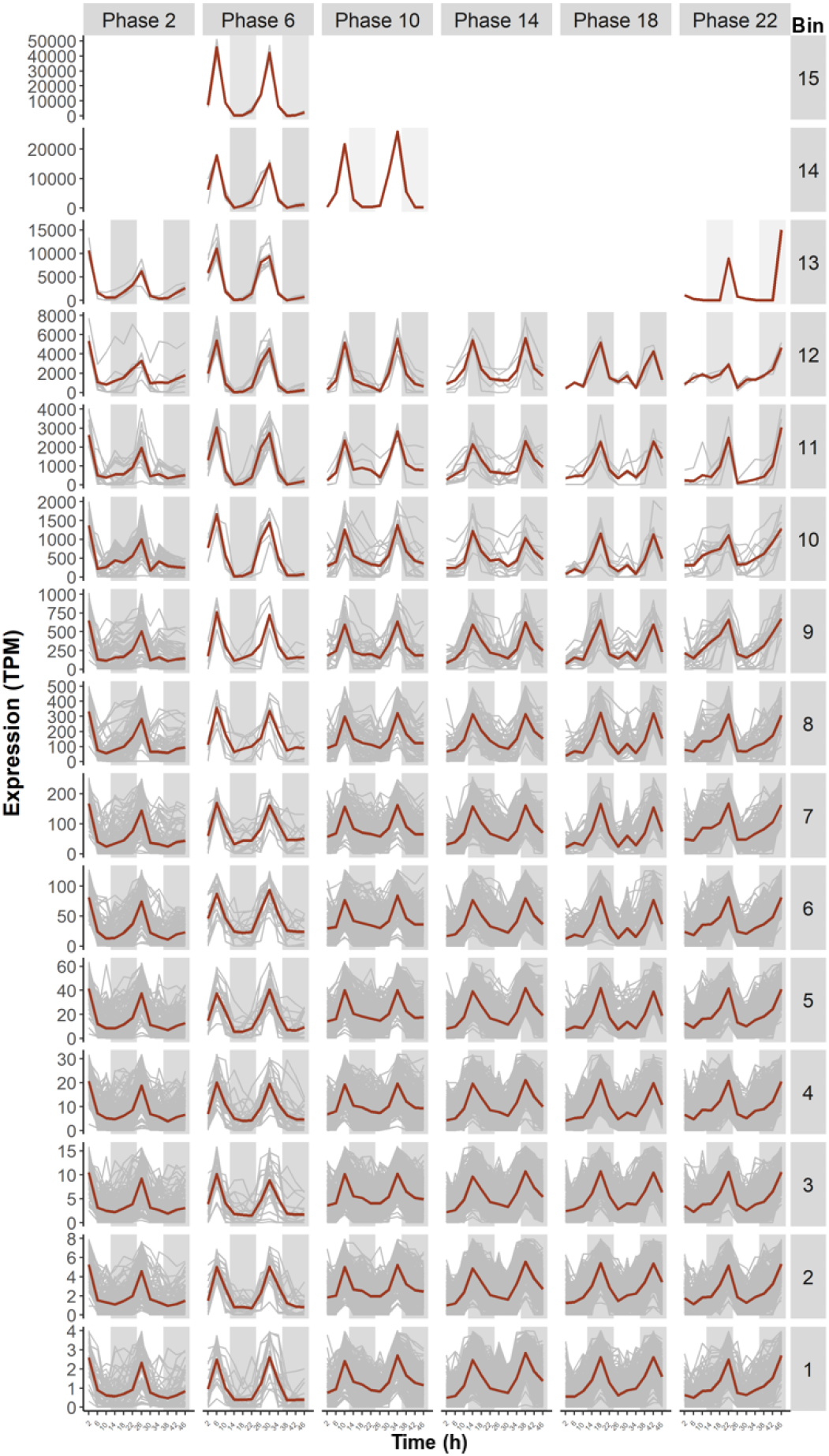
Expression of 24-h period rhythmic genes after classification by phase. Non-normalized transcripts per million (TPM) over the diurnal time series were plotted for 24-h period genes classified into phases, shown by horizontal panels (phases are indicated in grey boxes at the top). To improve the visibility of rhythmic expression over the large range of expression levels, genes were subdivided (binned) according to their maximal expression level, shown as vertical panels. Genes were binned by the floor function of their log2 maximal expression level, which is indicated in the grey boxes on the right. The expression of each gene in each class is shown by a grey line, while the red lines show the average TPM for all genes in this class.

**Supplementary Figure 2:**
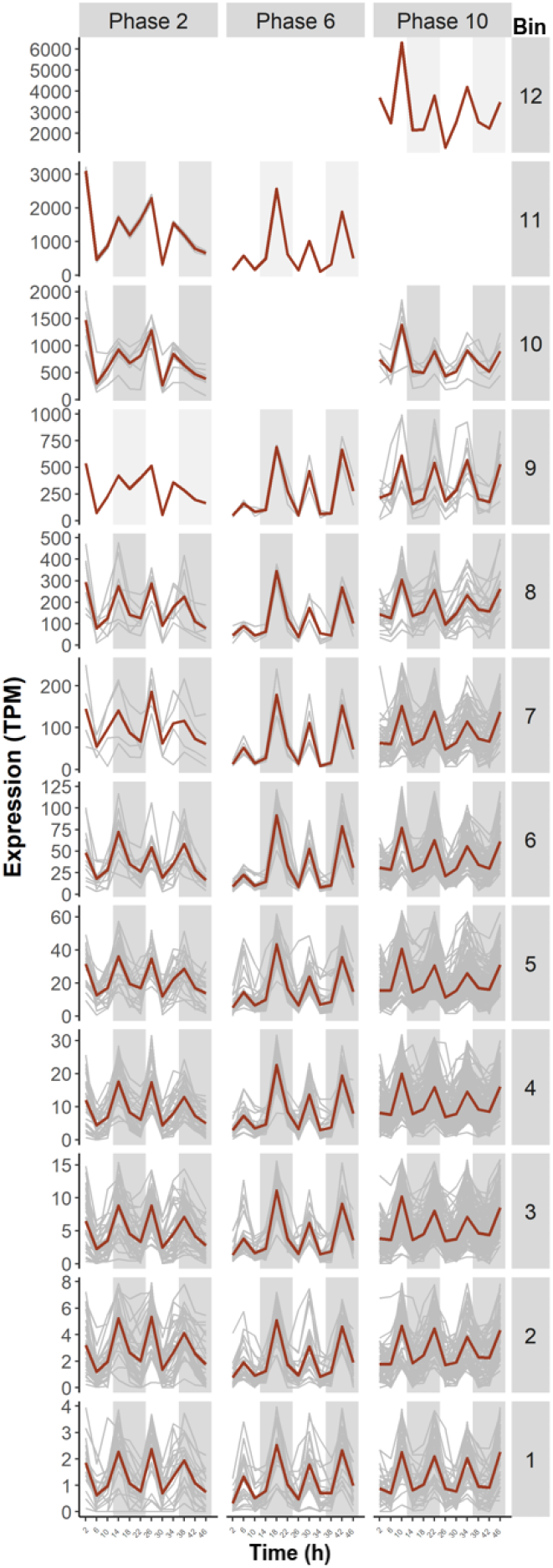
Expression of 12-h period rhythmic genes after classification by phase. Non-normalized transcripts per million (TPM) over the diurnal time series were plotted for 12-h period genes classified into phases, shown by horizontal panels (phases are indicated in grey boxes at the top). To improve the visibility of rhythmic expression over the large range of expression levels, genes were subdivided (binned) according to their maximal expression level, shown as vertical panels. Genes were binned by the floor function of their log2 maximal expression level, which is indicated in the grey boxes on the right. The expression of each gene in each class is shown by a grey line, while the red lines show the average TPM for all genes in this class.

**Supplementary Figure 3:**
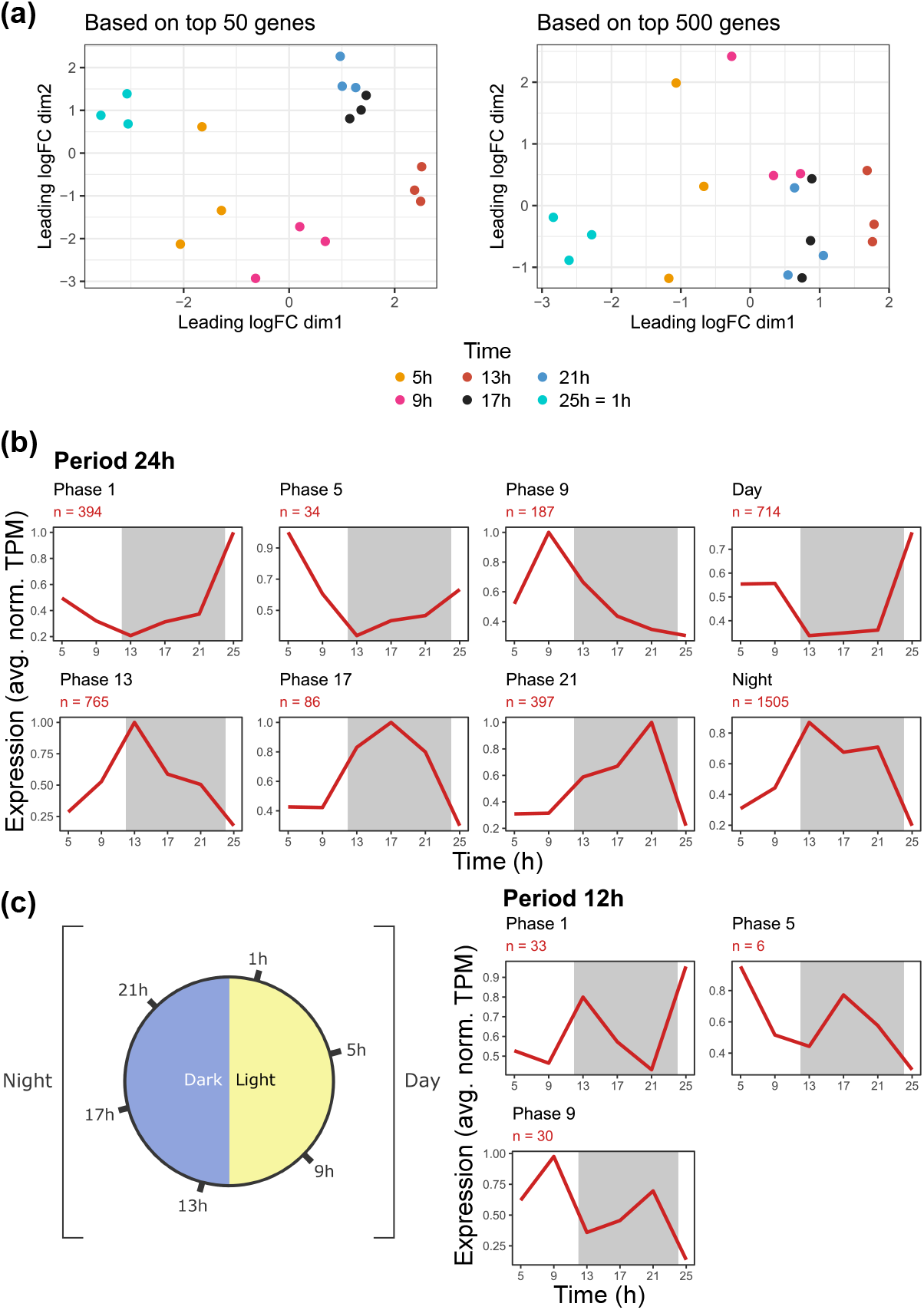
Overview of the nutrient-replete diurnal gene expression experiment from Smith et al. (2016) and classification of rhythmic *Phaeodactylum tricornutum* genes. **(a)** Multidimensional scaling (MDS) plot for the top 50 and 500 most differential genes of the 24-h diurnal time series. The top 50 was introduced as the top 500 MDS plot did not show a clear clustering with regard to time as was observed for *S. robusta*. **(b)** Average expression of gene sets of protein-coding genes for 24-h period (top) and 12-h period (bottom right) genes. Gene sets were defined based on EmpJTK period and phase prediction, as well as the position of the maximum in each day. Expression in transcripts per million (TPM) was normalized to a maximum of 1, and “n” represents the number of genes in each gene set. **(c)** schematic representation of the experimental setup and classification. The blue and yellow shaded halves represent the dark and light part of the 12h/12h cycle. Genes were classified in clusters for each phase (peak expression) as well as broader categories “Day” and “Night”, as indicated on the figure.

**Supplementary Figure 4:**
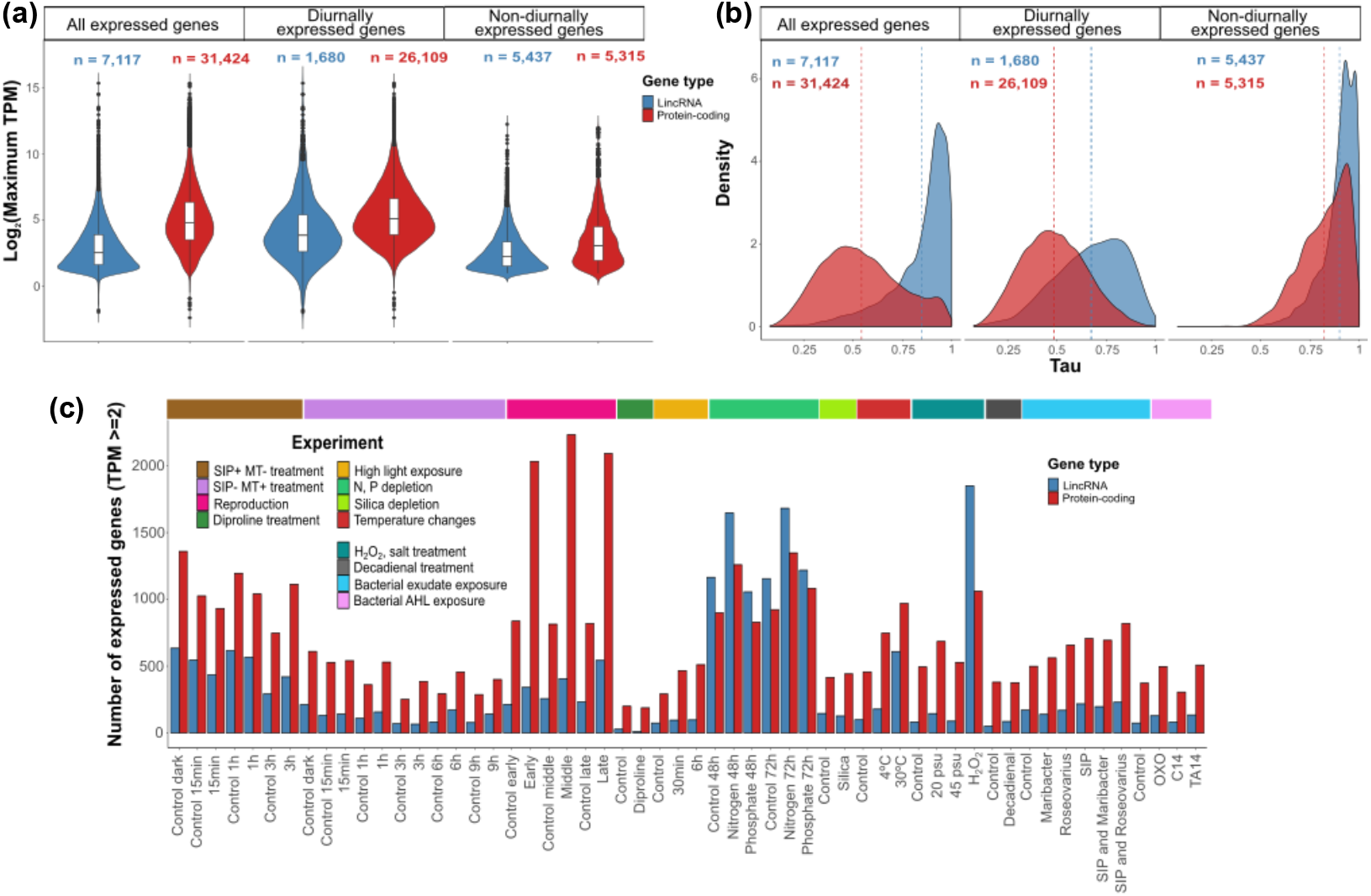
Overview of gene expression in experimental conditions outside the diurnal regime. Adaptor-trimmed reads from publicly available *S. robusta* transcriptomes [33] were mapped to the longest gene isoform using Salmon v1.1 (indexing using –keepDuplicates -k 31, quantification using –seqBias –gcBias -l A parameters) [124]. TPM refers to Transcripts Per Million. **(a)** Comparison of maximum TPM distribution of protein-coding and long intergenic non-coding RNAs (lincRNAs) in the *S. robusta* expression atlas without diel samples. Different sets of genes are introduced based on expression patterns: genes that are expressed in diurnal or other conditions (“All expressed genes”), the subset of genes that was expressed in the diurnal cycle (“Diurnally expressed genes”), and genes not expressed in the diurnal cycle, but expressed in other conditions (“Non-diurnally expressed genes”). The violin plot shows the kernel density estimation of the data. The line that divides the white box into two parts indicates the median of the data. The lower end of the box shows the Q1 quartile while the upper end the Q3 quartile. The lower whisker extends from the hinge to Q1-1.5xIQR while the upper whisker extends from the hinge to Q3+1.5xIQR. Data beyond the whiskers are potential outliers and are plotted as individual dots. **(b)** Comparison of Tau distribution of protein-coding genes and lincRNAs for different gene expression groups. Density refers to the kernel density estimate while Tau refers to the condition-specific expression metric [52]. TPM values were log2-transformed prior to Tau calculation. **(c)** Barplot showing the number of protein-coding and lincRNA genes expressed in non-diurnal conditions (control + treatment), including sexual reproduction, abiotic stress and bacteria-interaction experiments. Only non-diurnal expressed genes are included in this panel (5,315 protein-coding and 5,437 lincRNA genes).

**Supplementary Figure 5:**
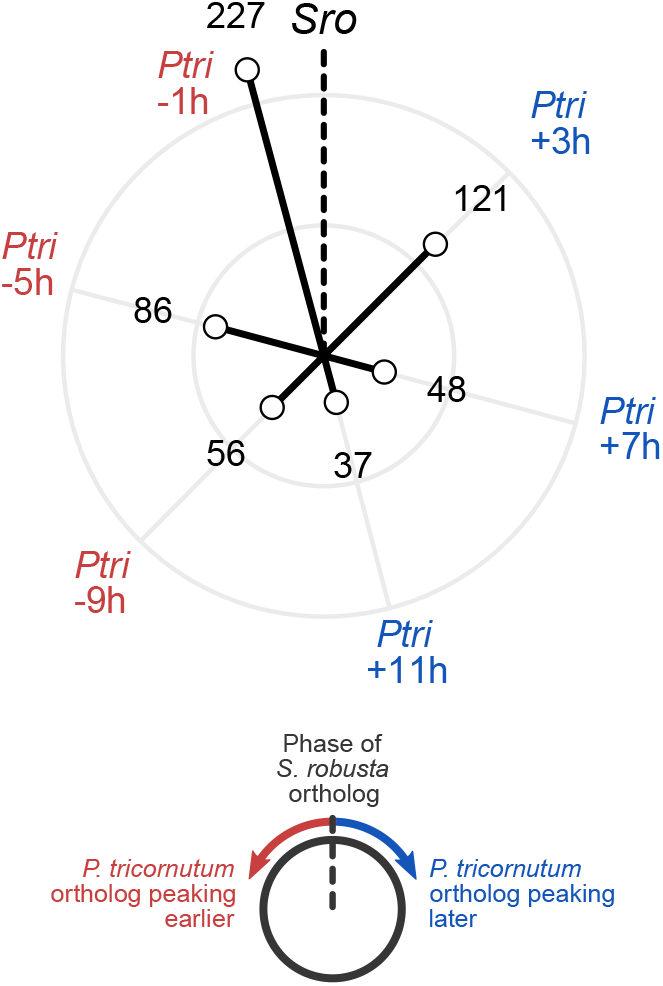
Comparative analysis of diel expression in *S. robusta* and *P. tricornutum*. Circular bar chart showing the phase of 575 *P. tricornutum* (“Ptri”) rhythmic genes compared to their one-to-one *S. robusta* (“Sro”) orthologs. The phase of the reference *S. robusta* ortholog is indicated with a dotted line, while the shift in phase of the corresponding *P. tricornutum* ortholog is shown on the outside of the circle. Each dataset consists of six time points per cycle in 4-h intervals, limiting the number of possible phase shifts to six. Since time series of *S. robusta* and *P. tricornutum* are offset by 1h, phase shifts of −1h are the most abundant and represent “conserved phasing”. The number of genes for each phase shift is shown by the length of each bar and is printed in black numbers. Shifts in which the *P. tricornutum* gene is peaking relatively earlier or later than its *S. robusta* counterpart are indicated in red or blue respectively.

**Supplementary Figure 6:**
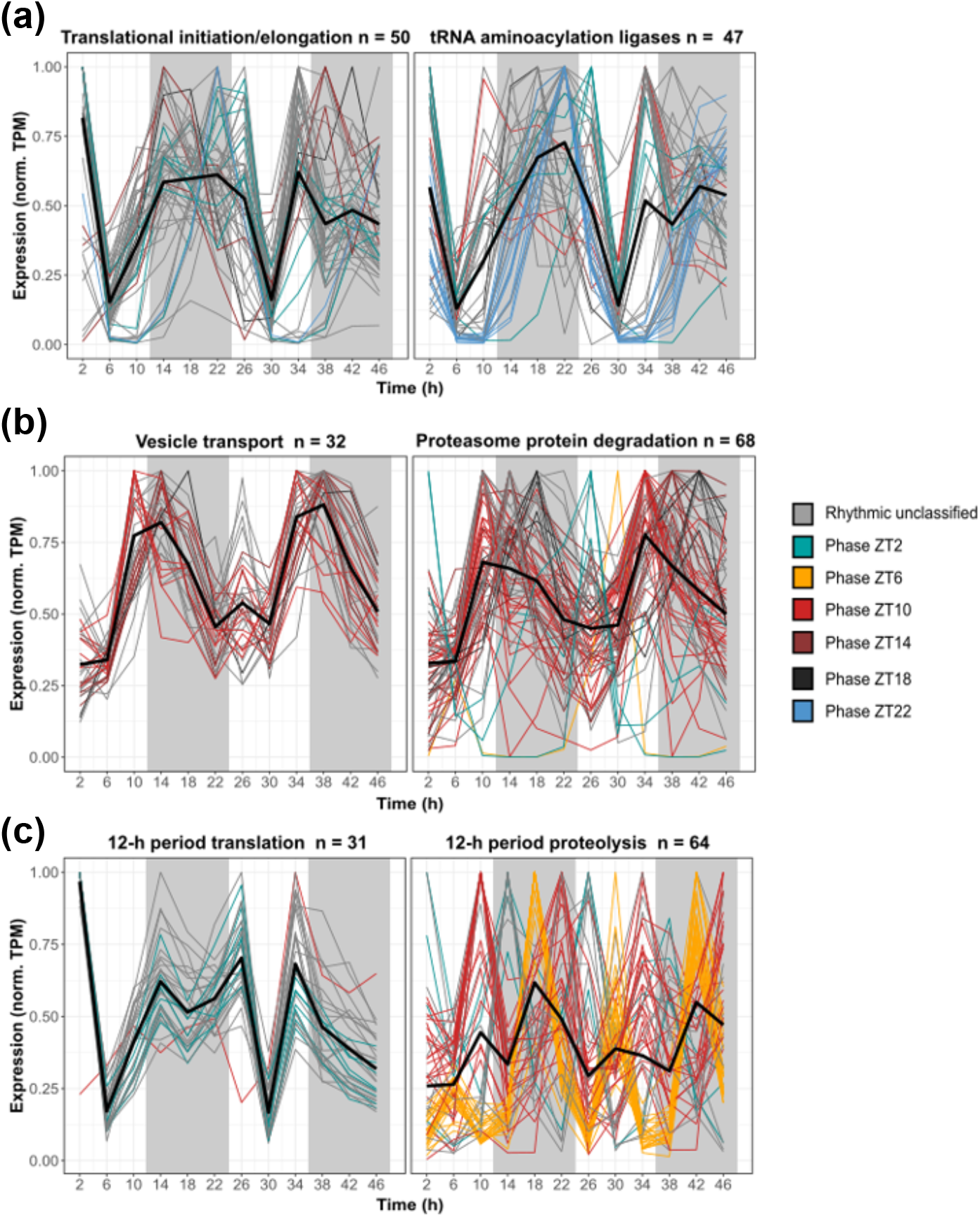
Overview of rhythmicity in protein turnover gene expression in *S. robusta*. Line plots showing normalized expression (transcripts per million, TPM) over time for different gene groups. Lines are coloured based on the gene phase, whether the phase is not consistent (unclassified, grey) or consistent (green = ZT2, yellow = ZT6, red = ZT10, brown = ZT14, black = ZT18, blue = ZT22). **(a)** All rhythmic genes (following a 24-h or 12-h period) annotated with “translational initiation” (GO:0006413) or “translational elongation” (GO:0006414) (left) and “tRNA aminoacylation for protein translation” (GO:0006418) (right). **(b)** All rhythmic genes (following a 24-h or 12-h period) annotated with “Golgi vesicle transport” (GO:0048193) or “retrograde transport, endosome to Golgi” (GO:0042147) (left) and “ubiquitin-dependent protein catabolic process” (GO:0006511) (right). **(c)** All 12-h periodical rhythmic genes annotated with “translation” (GO:0006440) (left) or “proteolysis” (GO:0006508) (right).

**Supplementary Figure 7:**
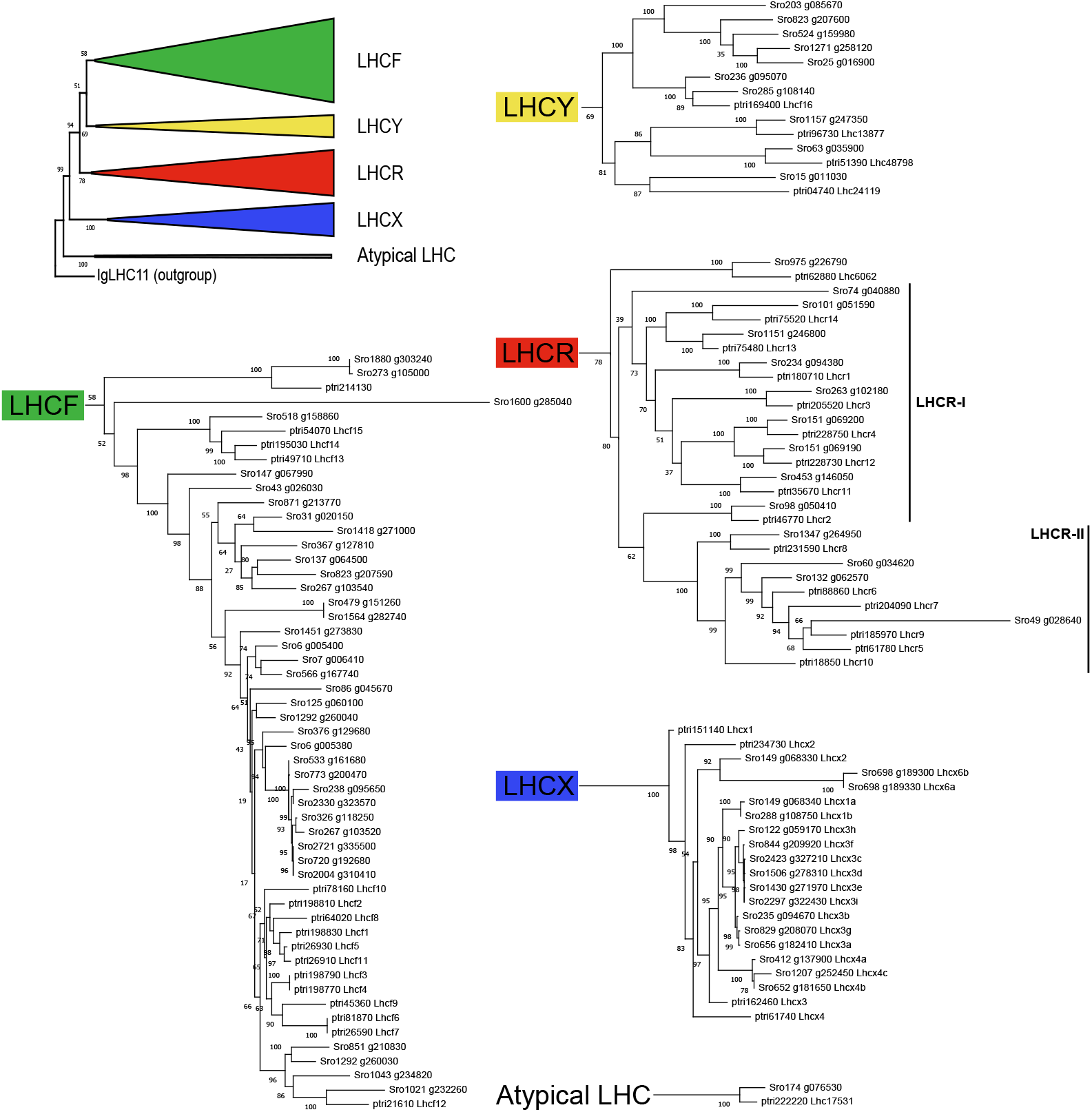
Maximum likelihood phylogenetic tree of the *S. robusta* and *P. tricornutum* LHC superfamily. Numbers indicate bootstrap support values at each branch. To improve readability, the full phylogenetic tree was split up in clades (LHCF, LHCX, LHCR and LHCY, indicated at the root of each subtree), the topology of major clades with respect to their outgroup (IgLHC11) is shown in the top left.

**Supplementary Figure 8:**
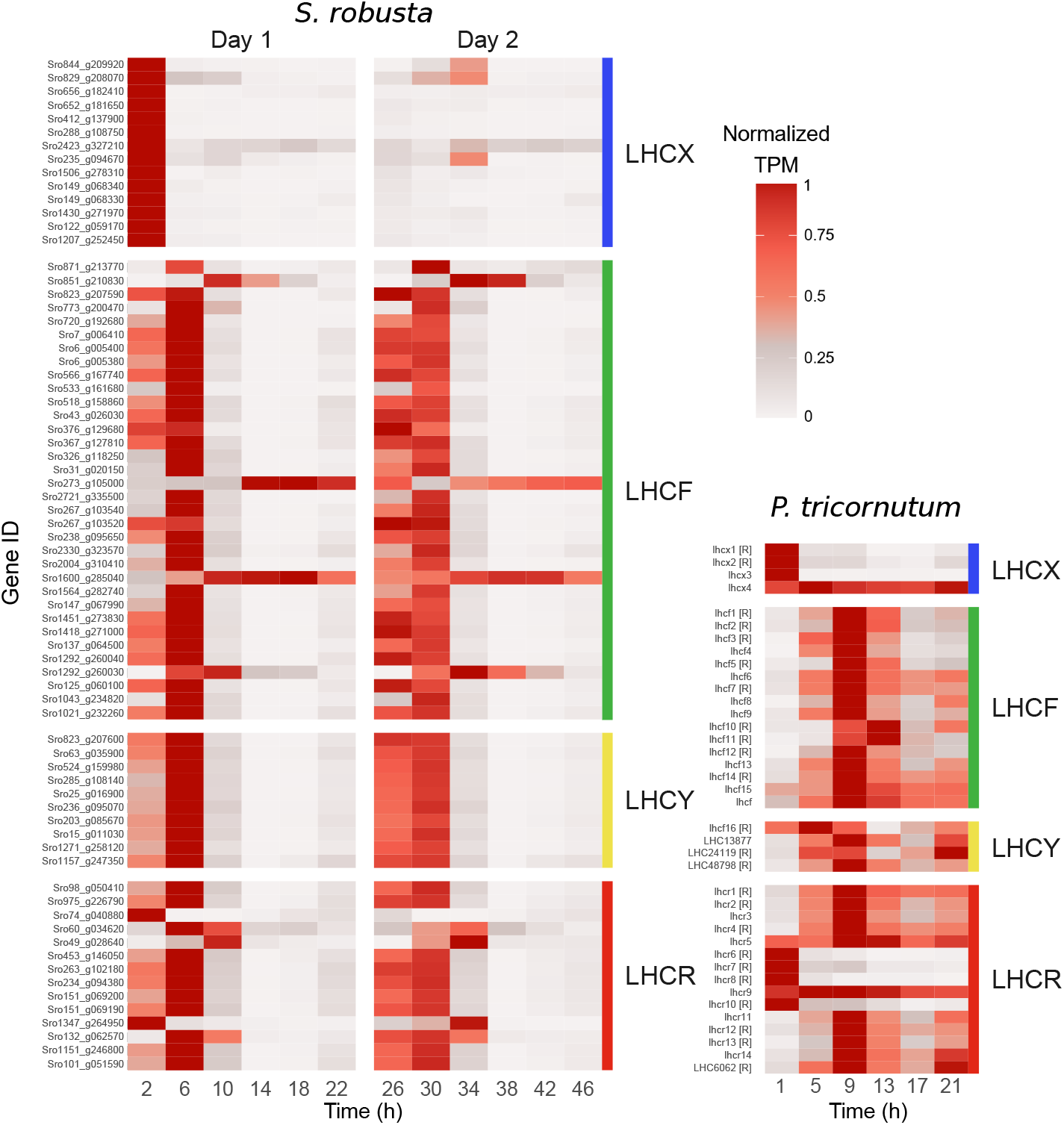
Heatmap of the expression of light harvesting complex proteins (LHC) in *S. robusta* and *P. tricornutum* [10] over the diel cycle. Expression data was plotted as average normalized TPM (transcripts per million) for each time point, ranging from no expression (grey) to high expression (dark red). The x-axis represents time (in hours) and the y-axis contains LHC genes. All expressed *S. robusta* LHC were significantly rhythmic, while [R] indicates significantly rhythmic genes for *P. tricornutum*. Vertical panels denote the four categories of LHC proteins present in diatoms.

**Supplementary Figure 9:**
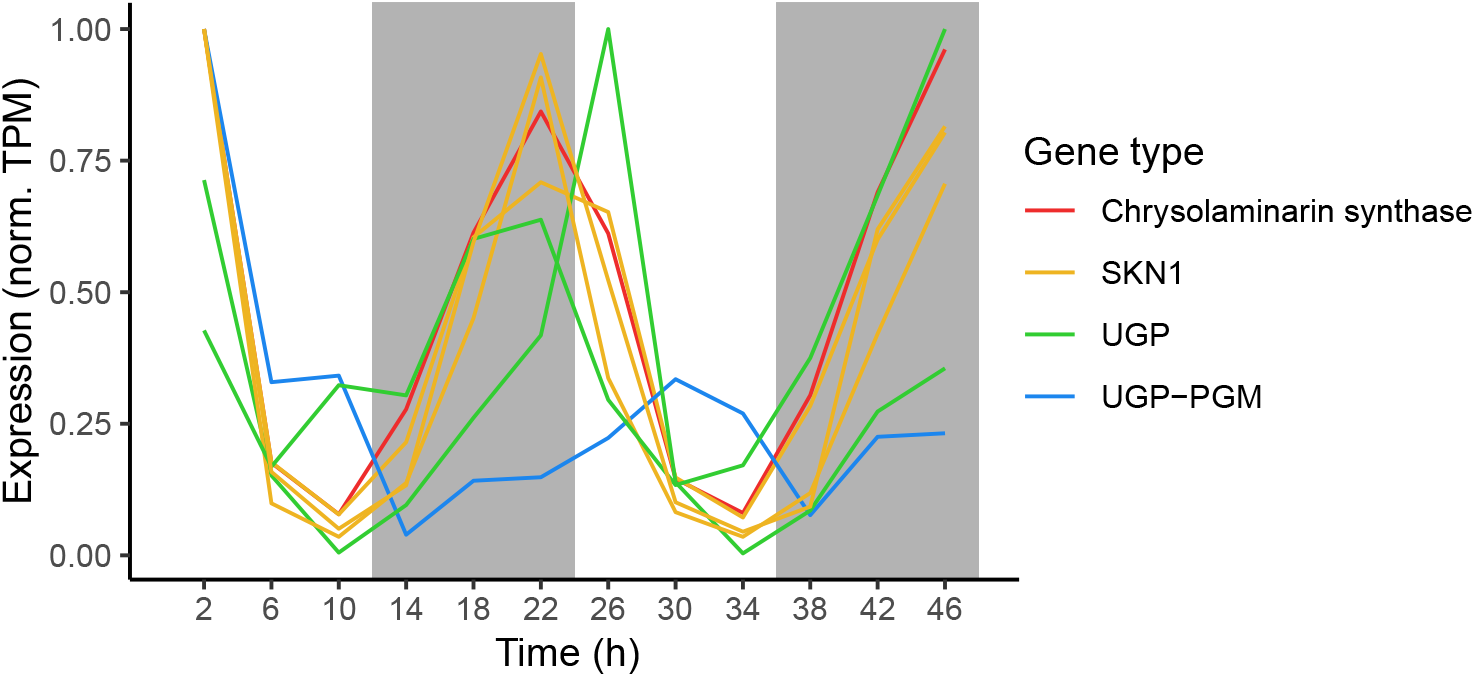
Diurnal expression of genes putatively involved in the synthesis of the storage carbohydrate chrysolaminarin. Expression for each gene is shown as a line connecting average normalized TPM (transcripts per million) over time. Grey shaded background boxes indicate periods of dark in the 12/12 day/night experiment, while the colour of each line represents the enzyme it encodes. SKN1 = beta-glucan synthesis-associated protein Suppressor of Kre6 Null, UGP = UDP-glucose pyrophosphorylase, UGP-PGM = UDP-glucose pyrophosphorylase/phosphoglucomutase.

**Supplementary Figure 10:**
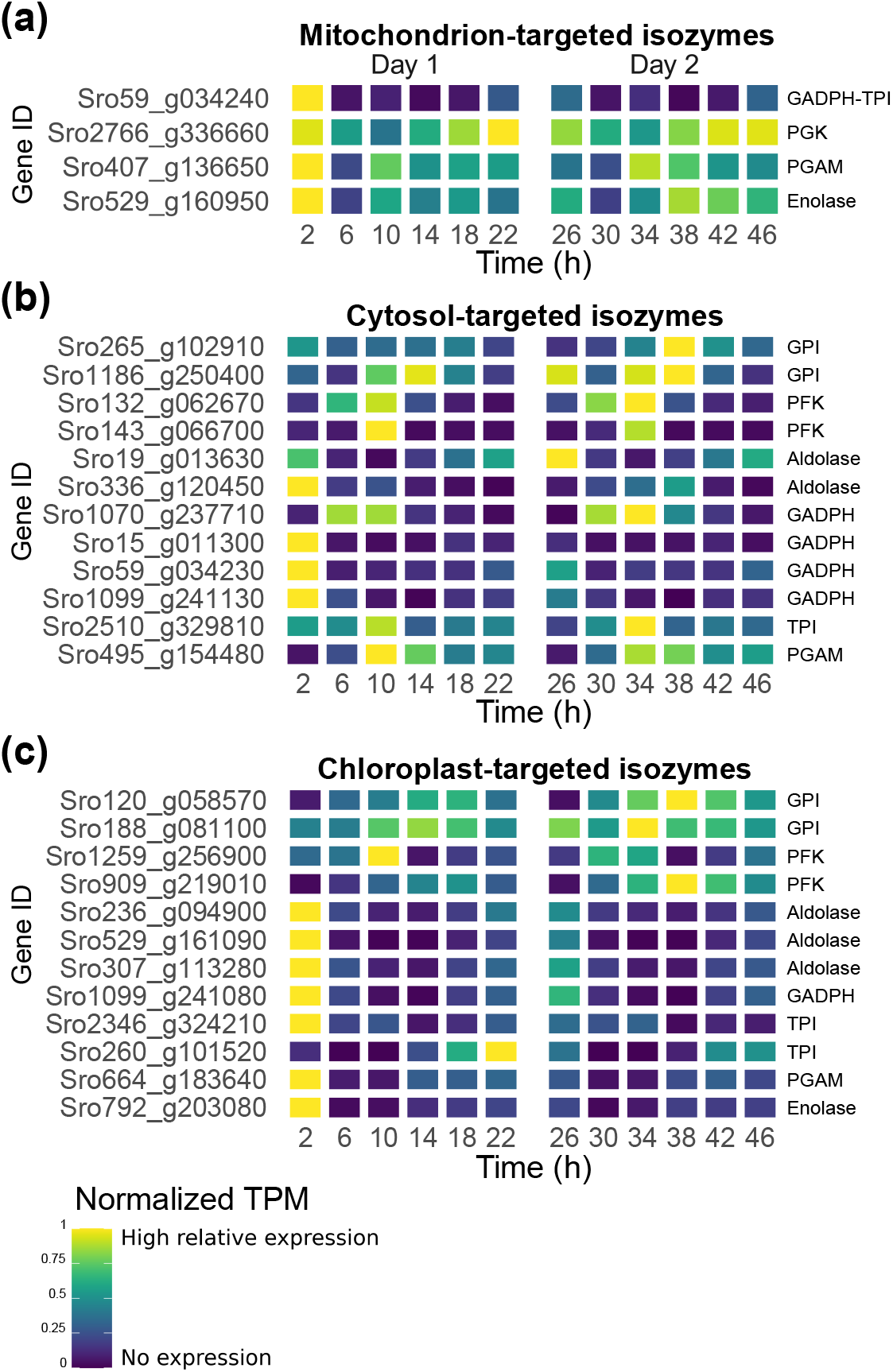
Heatmap of gene expression of rhythmic glycolytic isozymes. Heatmaps show the average normalized expression over two diurnal cycles of isozymes of the glycolysis/gluconeogenesis pathway, that are either predicted to be targeted to the mitochondrion **(a)**, the cytosol **(b)** or the chloroplast **(c)**. Average expression was calculated by normalizing transcripts per million (TPM) to a maximum of 1 over the time series. The short name of the enzyme is shown on the right side of the figure.

**Supplementary Figure 11:**
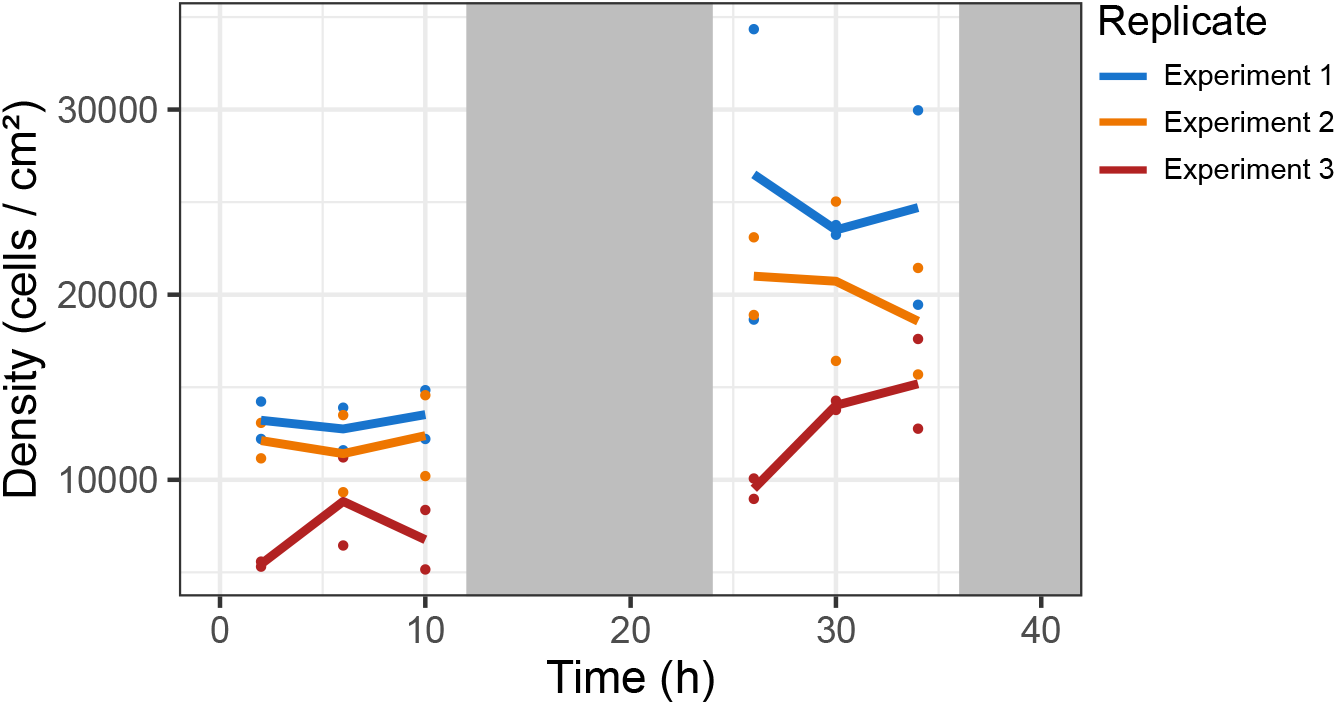
Culture density during the diurnal experiment. Density of *S. robusta* cultures during the three experimental repeats of the diurnal experiment (indicated by three different colours). Density of cells on the flask surface (cells / cm ^2^) was determined by manual counting of cells in microscopic pictures obtained during the experiment. Points represent single measurements of technical replicates, while lines connect the average for each experimental repeat (biological replicate) over time. During the night, no pictures were taken from the flasks to avoid exposure to light. Grey shaded background boxes indicate periods of darkness in the 12/12 day/night experiment.

**Supplementary Figure 12:**
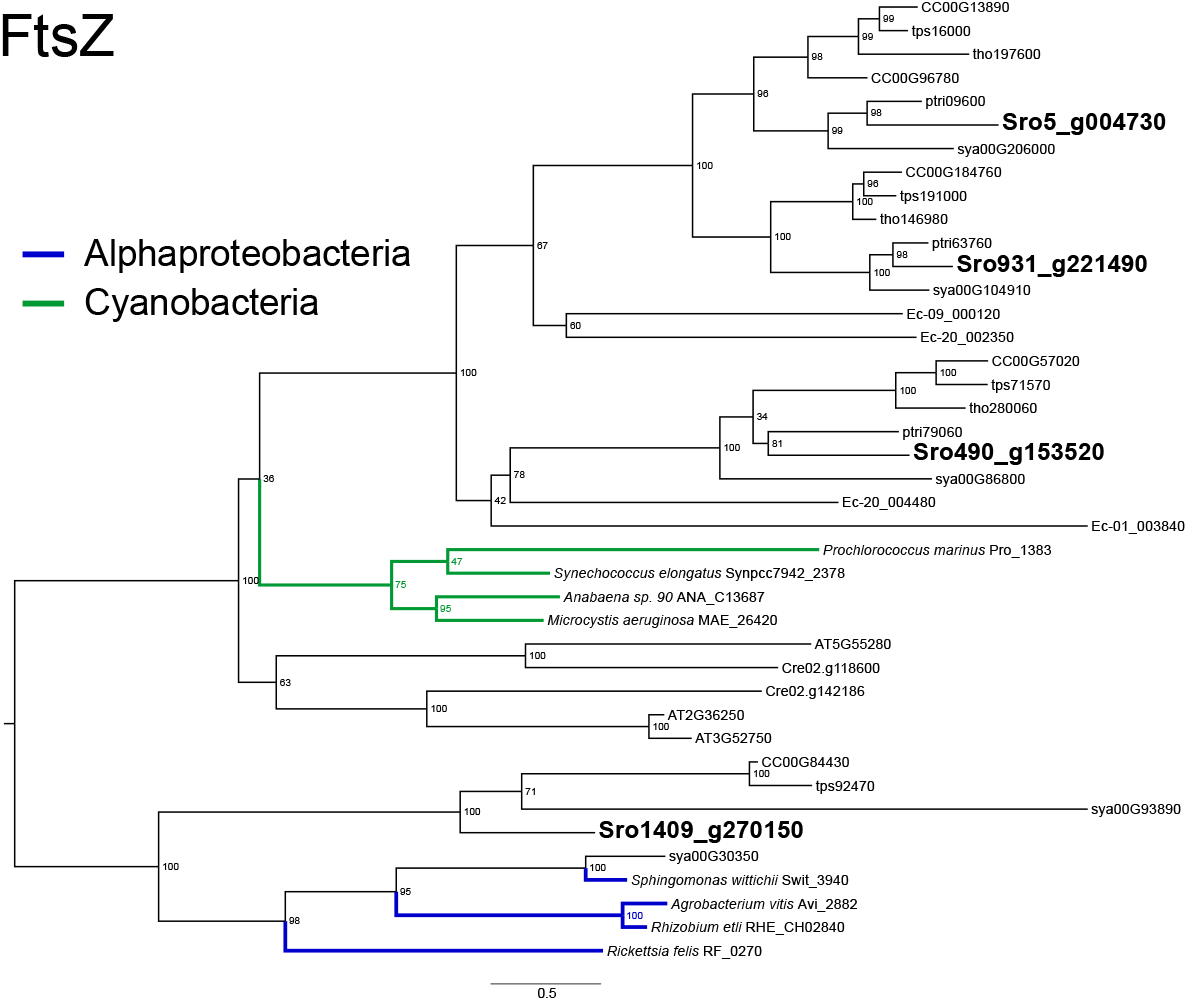
Maximum likelihood phylogenetic tree of FtsZ proteins. Midpoint-rooted maximum likelihood phylogenetic tree of FtsZ genes in selected eukaryotic species, as well as homologs from Alphaproteobacteria (blue) and Cyanobacteria (green). *S. robusta* genes were printed in bold and enlarged, and were assigned to be of mitochondrial or chloroplastic origin based on their clustering pattern with bacterial homologs. Numbers at each branch point are bootstrap support values. Eukaryotic gene IDs refer to PLAZA Diatoms identifiers.

**Supplementary Figure 13:**
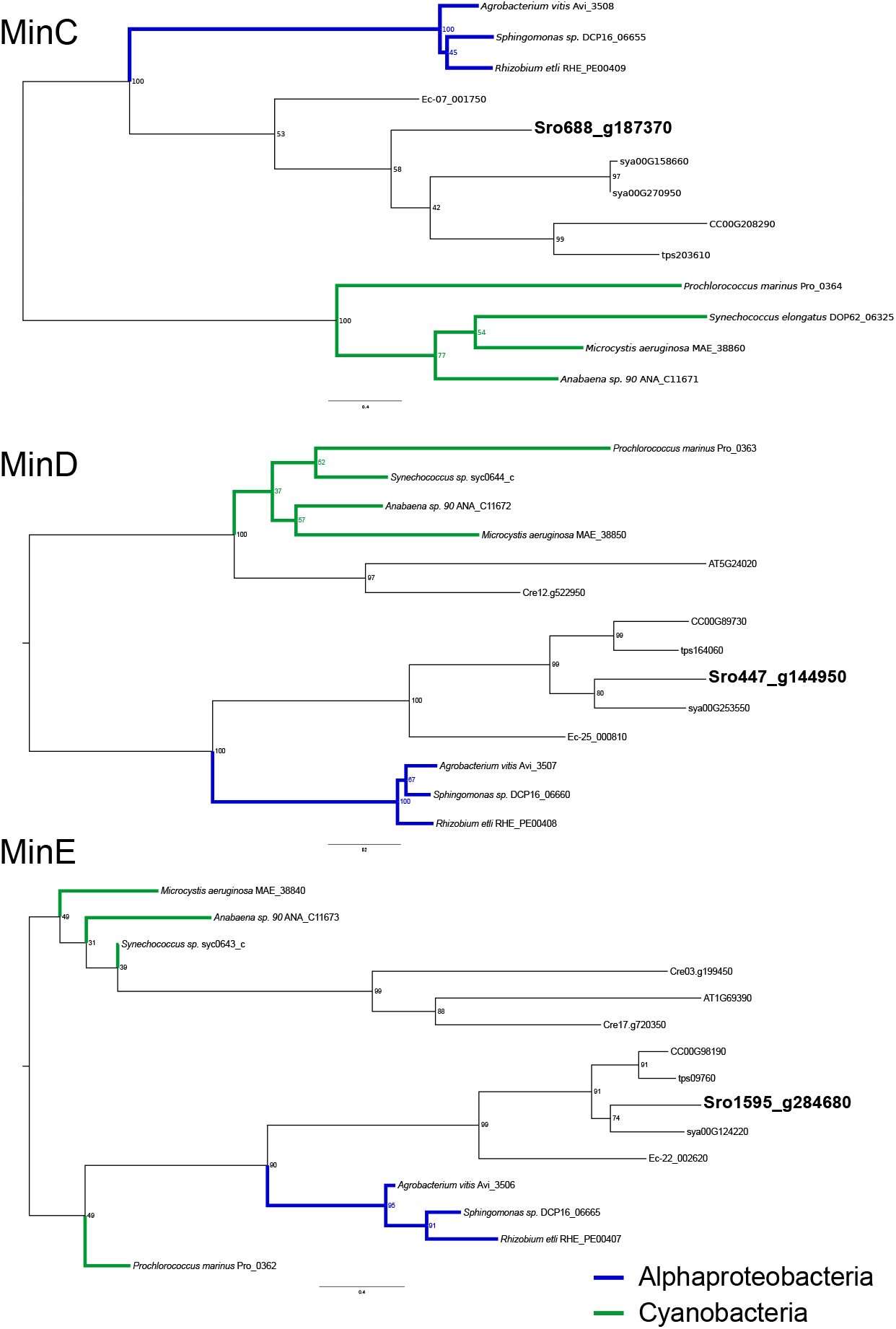
Maximum likelihood phylogenetic tree of Min-system proteins. Midpoint-rooted maximum likelihood phylogenetic tree of genes from the Min system (MinC, MinD and MinE) in selected eukaryotic species, as well as homologs from Alphaproteobacteria (blue) and Cyanobacteria (green). *S. robusta* genes were printed in bold and enlarged, and were assigned to be of mitochondrial or chloroplastic origin based on their clustering pattern with bacterial homologs. Numbers at each branch point are bootstrap support values. Eukaryotic gene IDs refer to PLAZA Diatoms identifiers.

**Supplementary Figure 14:**
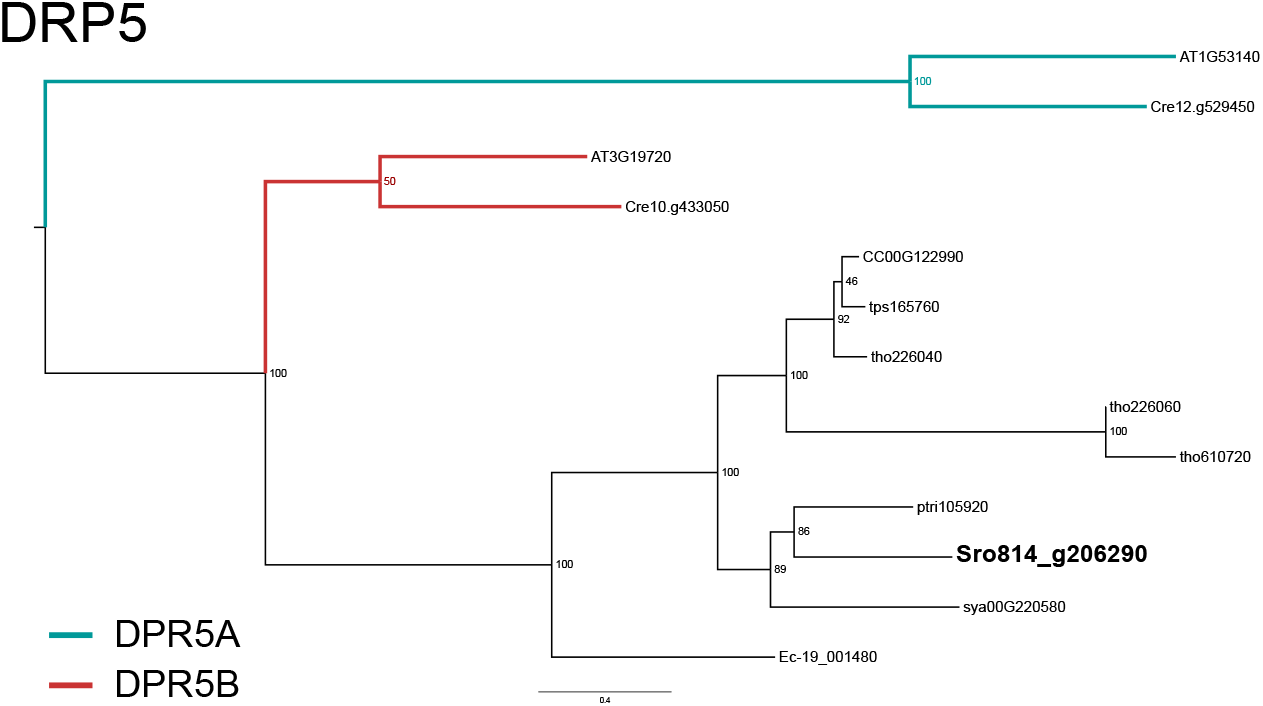
Maximum likelihood phylogenetic tree of DRP5A and DRP5B proteins. Midpoint-rooted maximum likelihood phylogenetic tree of dynamin-related GTPase DRP5 genes in selected eukaryotic species. Characterized DRP5A or DRP5B in plants (*Arabidopsis thaliana* and *Chlamydomonas reinhardtii*) are indicated with blue and red branches respectively. The *S. robusta* DRP5 gene was printed in bold and enlarged, and was characterized as a DRP5B homolog based on its phylogenetic position. Numbers at each branch point are bootstrap support values.

**Supplementary Figure 15:**
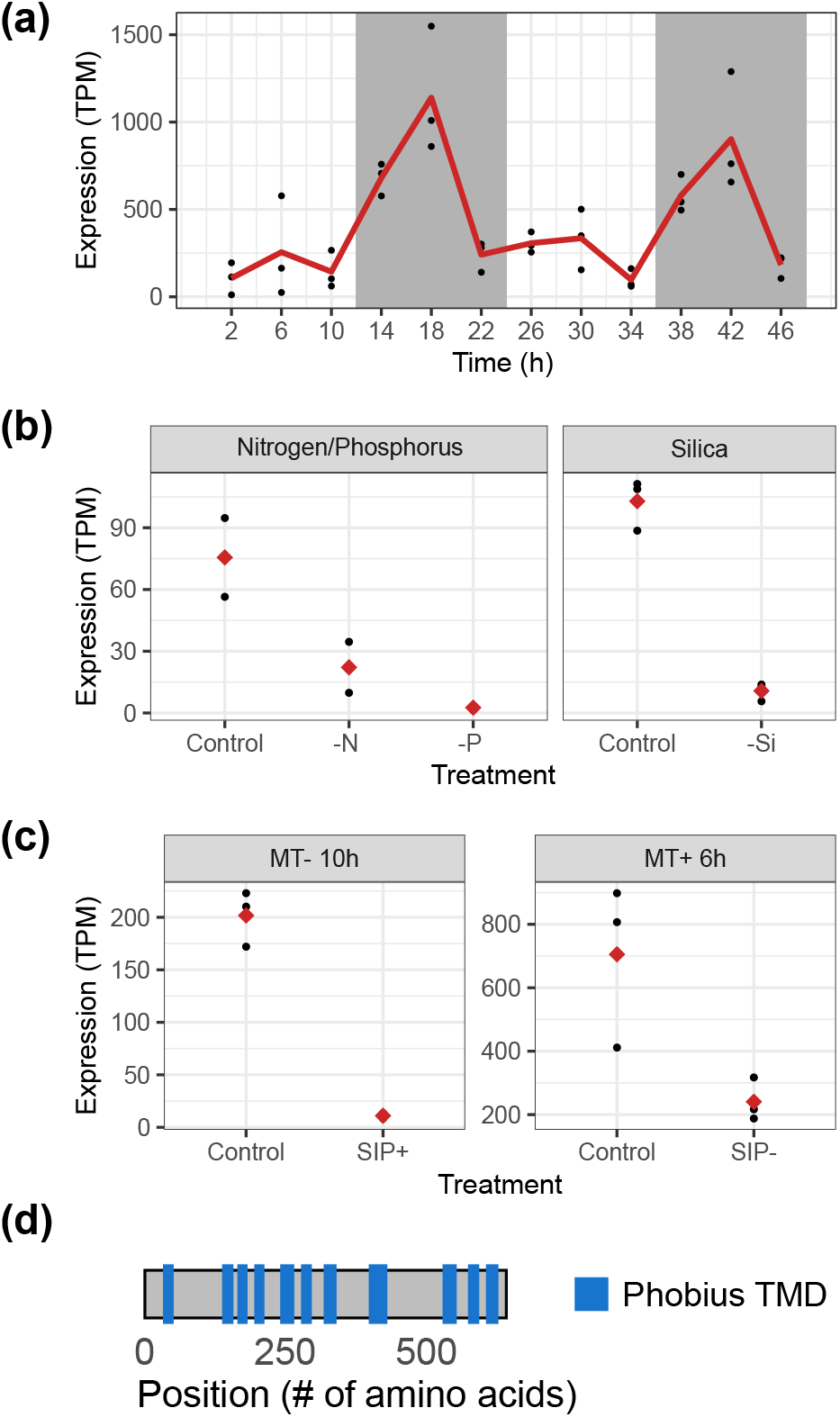
Expression and structure of the plant-type silicon efflux transporter-like gene Sro979_g227220 in *Seminavis robusta*. **(a)** Expression (transcripts per million, TPM) during the diurnal cycle is plotted in function of time. Points show single replicate measurements, while the red line connects the average at each time point. Grey shaded areas show the dark treated portions of the experiment. **(b)** expression in response to nutrient limitation, retrieved from the *S. robusta* expression atlas in Osuna-Cruz et al. (2020) [33]. Black points indicate individual measurements, while red diamonds indicate the average TPM for each treatment level. Control = nutrient replete conditions, -N = 72h of Nitrogen depletion, -P = 72h of Phosphorus depletion and -Si = 24h of Silicon depletion. **(c)** TPM with (SIP) and without (Control) addition of the cell cycle arresting sex inducing pheromone from the opposite mating type (MT) after 10h in MT- (left) and 6h in MT+ (right). Data retrieved from Bilcke et al. (2021) [32] and Cirri et al. (2019) [35]. **(d)** Position of the 11 transmembrane domains in the gene Sro979_g227220 as predicted by Phobius.

**Supplementary Figure 16:**
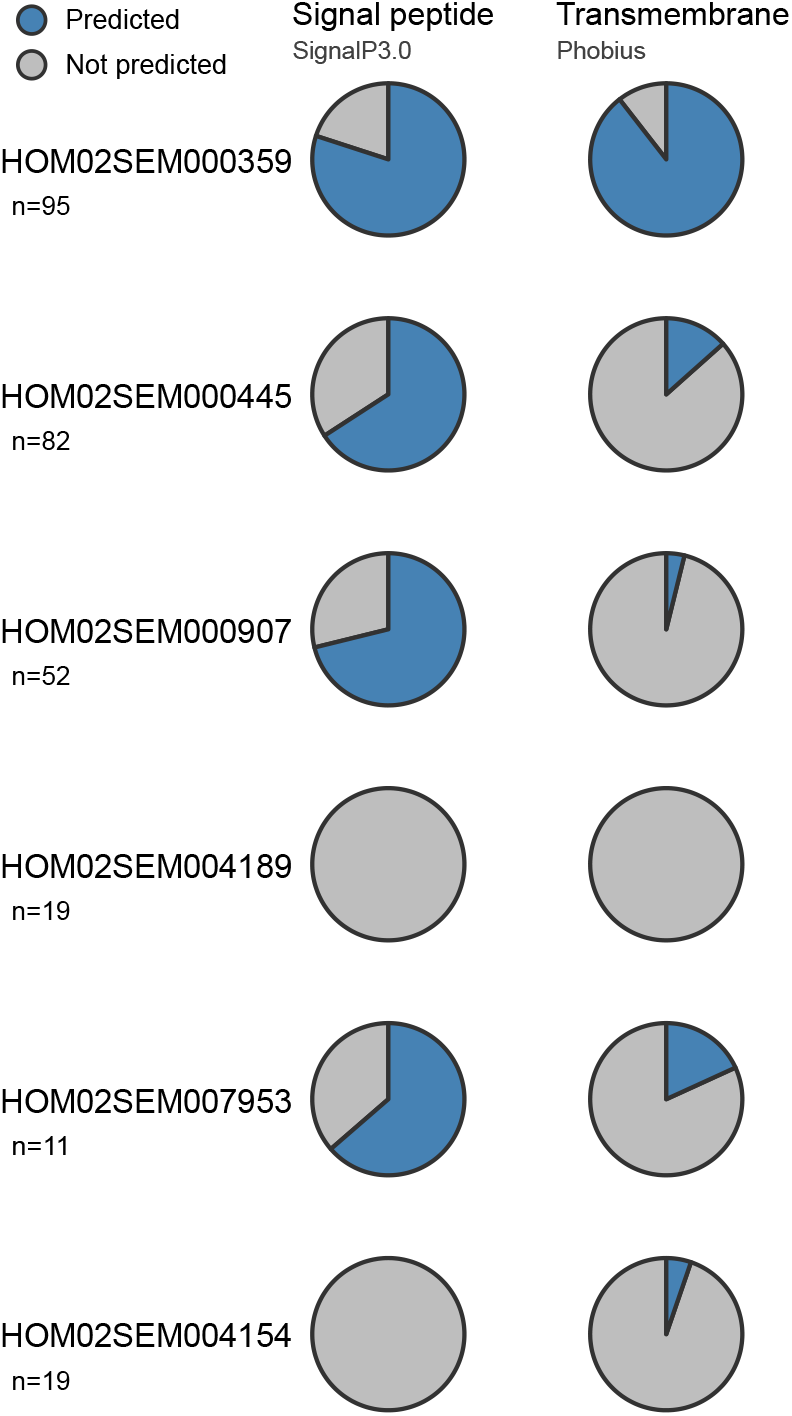
Signal peptide and transmembrane domain prediction of known and putative new diatom cell wall families. Pie charts show the proportion of genes in each gene family that have a predicted signal peptide or at least one transmembrane domain (blue). The PLAZA Diatoms gene family identifier and the total number of genes encoded in the 10 diatom genomes for each family (n) are given on the left.

**Supplementary Figure 17:**
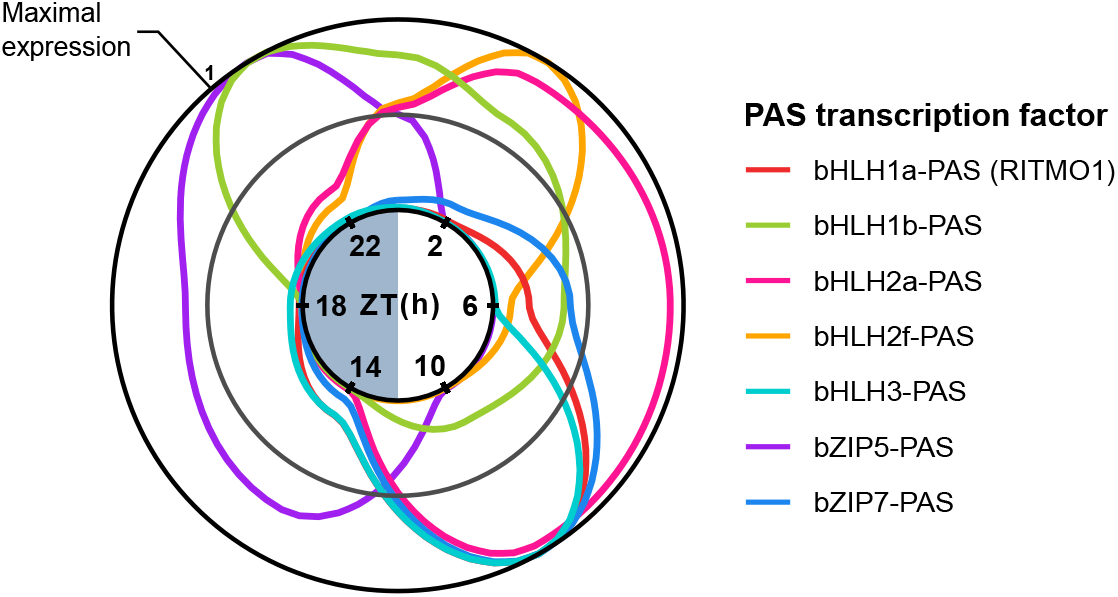
Expression peaks of potential circadian transcription factors containing a PAS domain cover the diurnal cycle. Expression in terms of normalized TPM (transcripts per million) averaged over replicates and days is shown for a selection of putative circadian PAS-containing transcription factors in *S. robusta* (some bHLH2-PAS homologs were omitted for clarity). Expression over Zeitgeber Time (ZT, hours since first illumination, see central circle) in hours was plotted using a Loess smoother connecting the average expression at each time point, after which the x-axis was circularized to show the continuous progression of the diurnal cycle. Expression of different genes is shown in a different coloured line.

**Supplementary Figure 18:**
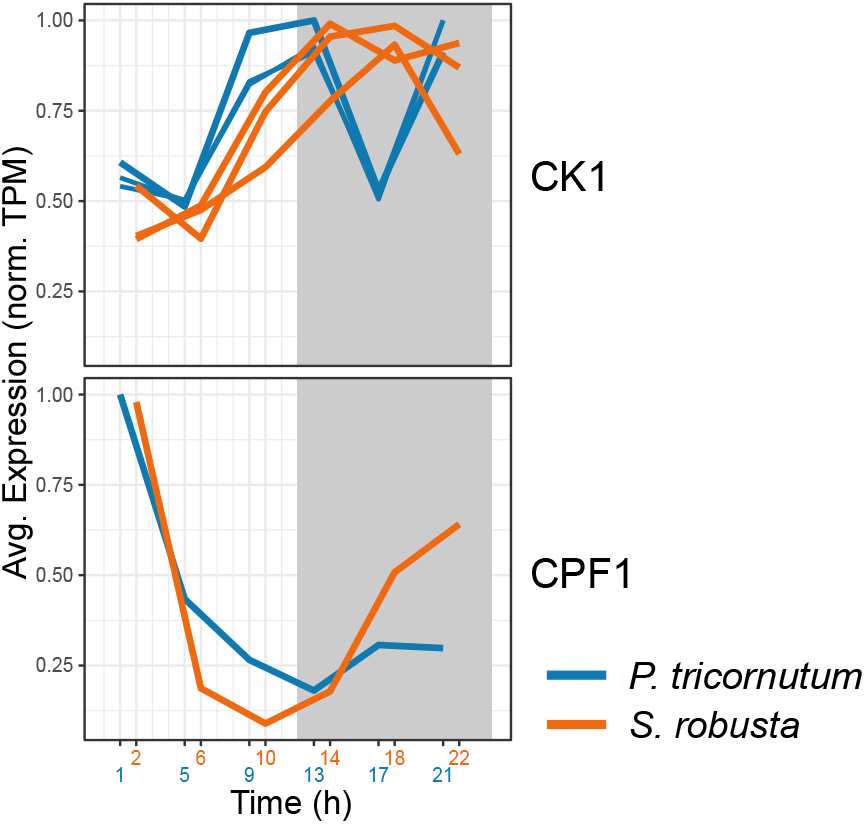
Expression of casein kinase 1 (CK1) and cryptochrome/photolyase family 1 (CPF1) genes. Normalized expression (transcripts per million, TPM) over time is shown for each candidate gene (S. *robusta* = orange, *P. tricornutum* = blue). To allow comparison with the 1-day *P. tricornutum* time series, *S. robusta* relative expression was averaged over both days and shown over the span of one day. The *P. tricornutum* time point 25h represents the equivalent 1h in these plots.

**Supplementary Figure 19:**
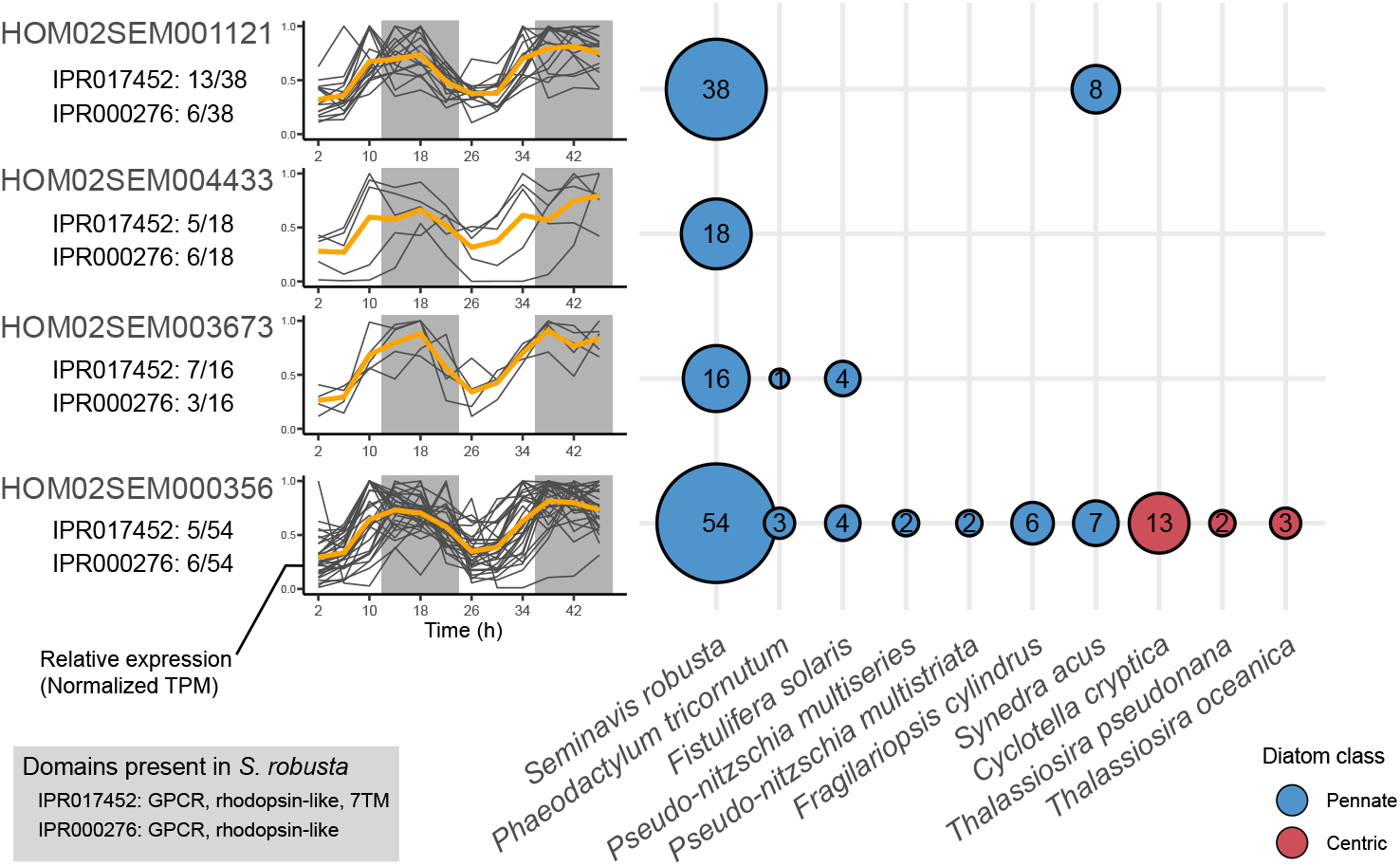
Expression and distribution of potential G-protein coupled receptor (GPCR) rhodopsin photoreceptors. Relative expression of genes belonging to four putative rhodopsin families during the diel cycle is plotted at the left side as TPM normalized for each gene to a maximum of 1 over the time series. Only significantly rhythmic genes with a 24-h period were plotted. Solid lines connect observations from the same gene over time, while the orange line connects the average relative expression for each family over the time course. At the right side, the presence of members of each family in the genome of 10 diatoms is shown, with circle radii and numbers representing the number of genes per species, and fill colours indicating if species belong to centric or pennate diatoms. Below the gene family ID, we have indicated the number of genes in the family that contain a predicted rhodopsin-like InterPro protein domain.

**Supplementary Figure 20:**
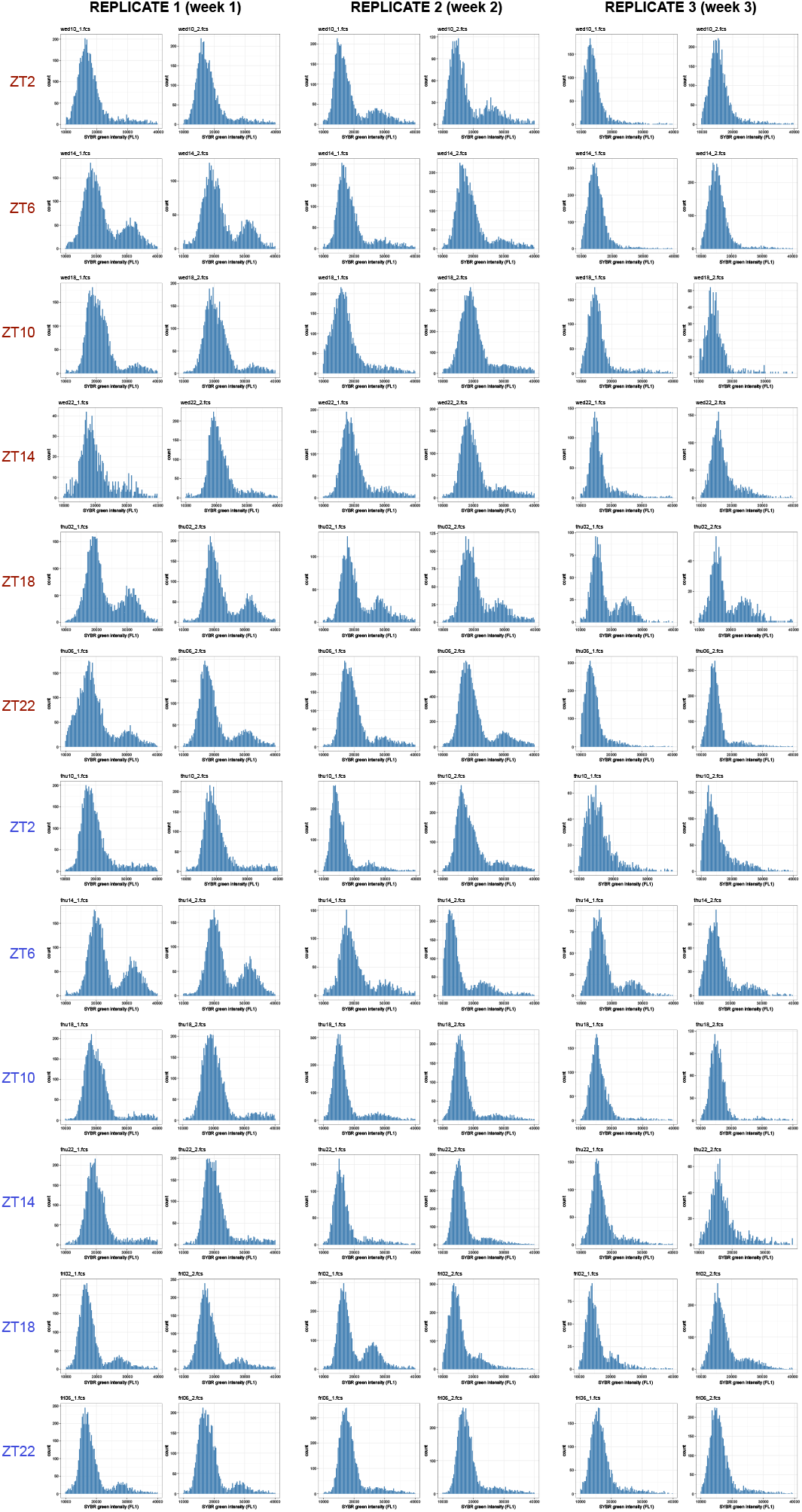
Histograms of flow cytometry during the diurnal cycle experiment. Histograms showing the frequency of SYBR green intensity (FL1) during the diurnal experiment. Two technical replicates were measured for each of the three experimental repeats (horizontal) and each time point (vertical, ZT = Zeitgeber Time, time in h since illumination). Time point colours indicate the day: red = first day, blue = second day.

## 9 Supplementary Tables

**Supplementary Table 1:** Classification of rhythmic genes in *S. robusta* (Sro) and *P. tricornutum* (Ptri). This table shows the absolute number of genes assigned to each cluster, as well as the proportion relative to all diurnally expressed genes between parentheses. Columns display the frequencies observed in the total set of expressed genes, the set of protein coding and long intergenic non-coding RNA (lincRNA) genes. The total number of annotated genes in the genome for each category is given in the column title (n =). The *P. tricornutum* results are based on a re-analysis of existing data from Smith et al. (2016) [10] and consist of only protein-coding genes.

| Classification cluster | Sro total<br>n = 43,371 | Sro protein-coding<br>n = 36,254 | Sro lincRNA<br>n = 7,117 | Ptri protein-coding<br>n = 12,072 |
| --- | --- | --- | --- | --- |
| Expressed genes in dataset | 27,789 (100%) | 26,109 (100%) | 1680 (100%) | 10,771 (100%) |
| Not significantly rhythmic | 3337 (12%) | 2781 (10,7%) | 556 (33,1%) | 8255 (76.6%) |
| Significantly rhythmic | 24,452 (88%) | 23,328 (89,3%) | 1124 (66,9%) | 2516 (23.4%) |
| *Period 24h* | 22,609 (81,4%) | 21,576 (82,6%) | 1033 (61,5%) | 2399 (22.3%) |
| No consistent phase | 11,279 (40,6%) | 10,625 (40,7%) | 654 (38,9%) | 536 (5%) |
| Early day (ZT2, ZT1) | 1113 (4%) | 1083 (4,1%) | 30 (1,8%) | 394 (3.7%) |
| Midday (ZT6, ZT5) | 220 (0,8%) | 213 (0,8%) | 7 (0,4%) | 34 (0.3%) |
| Late day (ZT10, ZT9) | 1978 (7,1%) | 1943 (7,4%) | 35 (2,1%) | 187 (1.7%) |
| Early night (ZT14, ZT13) | 3736 (13,4%) | 3588 (13,7%) | 148 (8,8%) | 765 (7.1%) |
| Midnight (ZT18, ZT17) | 2383 (8,6%) | 2298 (8,8%) | 85 (5,1%) | 86 (0.8%) |
| Late night (ZT22, ZT21) | 1900 (6,8%) | 1826 (7%) | 74 (4,4%) | 397 (3.7%) |
| Day | 1711 (6,2%) | 1662 (6,4%) | 49 (2,9%) | 714 (6.6%) |
| Night | 11,176 (40,2%) | 10,591 (40,6%) | 585 (34,8%) | 1505 (14%) |
| Dawn | 2222 (8%) | 2179 (8,3%) | 43 (2,6%) | NA |
| Dusk | 7369 (26,5%) | 7195 (27,6%) | 174 (10,4%) | NA |
| *Period 12h* | 1843 (6,6%) | 1752 (6,7%) | 91 (5,4%) | 117 (1.1%) |
| No consistent phase | 732 (2,6%) | 679 (2,6%) | 53 (3,2%) | 48 (0.4%) |
| Early day (ZT2, ZT1) | 184 (0,7%) | 171 (0,7%) | 13 (0,8%) | 33 (0.3%) |
| Midday (ZT6, ZT5) | 195 (0,7%) | 184 (0,7%) | 11 (0,7%) | 6 (0.1%) |
| Late day (ZT10, ZT9) | 732 (2,6%) | 718 (2,8%) | 14 (0,8%) | 30 (0.3%) |

**Supplementary Table 2:**
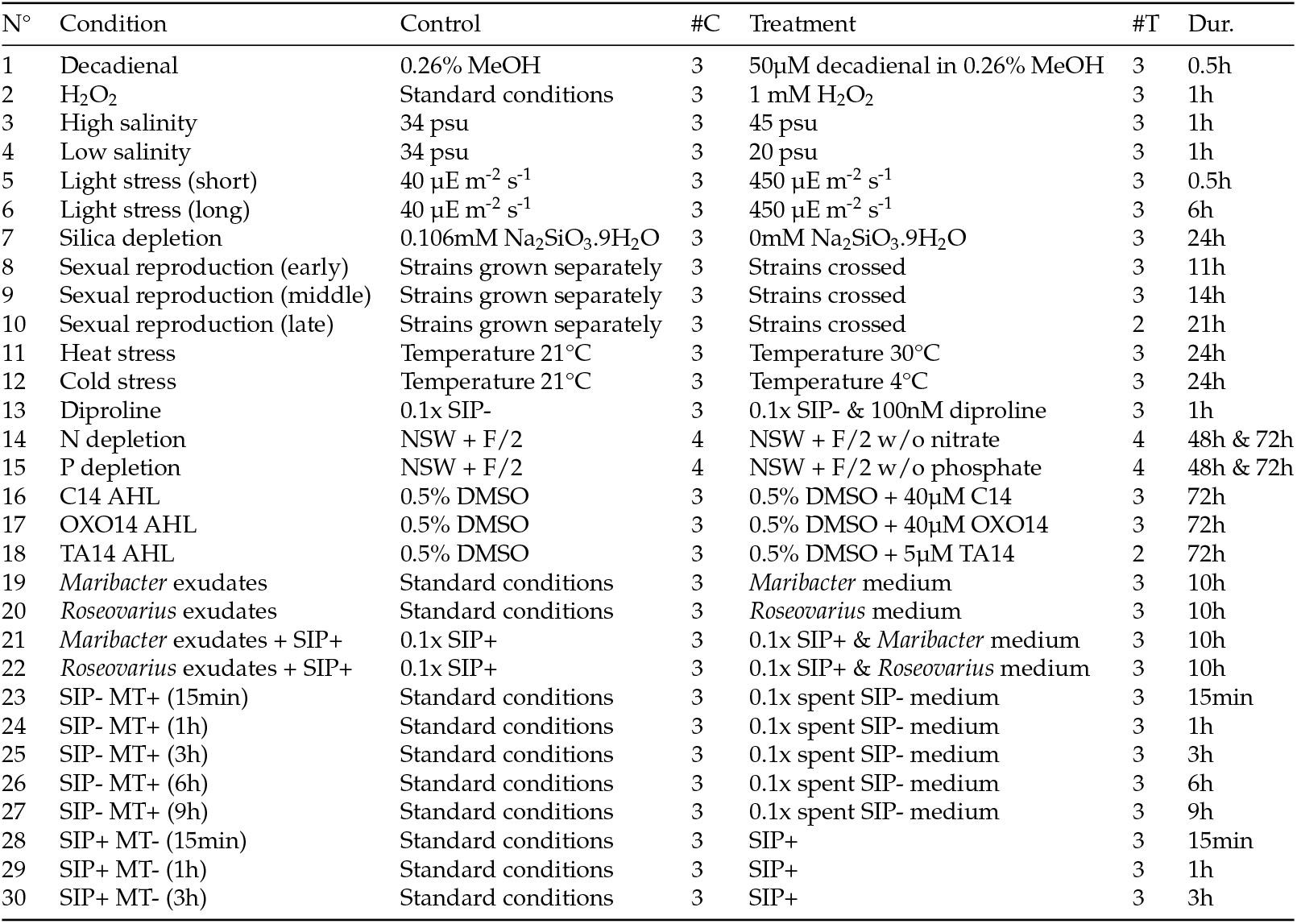
Overview of 30 conditions of the *S. robusta* gene expression atlas used in this paper. *S. robusta* cultures were subjected to 30 different treatments that range from abiotic stressors and biotic interactions to life cycle (sexual reproduction) stages. For each condition, replicates of control (untreated) and treated cultures were harvested and polyA-enriched mRNA was sequenced. Each condition represents one comparison (contrast) for which differential expressed genes were inferred. The day/night contrast was not included since it is superfluous to the full diurnal dataset presented in this work. #C: number of control replicates, #T number of treatment replicates. Dur.: duration of the treatment. AHL: N-acyl homoserine lactones and derivatives. NSW: natural sea water. SIP: sex inducing pheromone. RNA-seq reads were retrieved from Osuna-Cruz et al. (2020).

**Supplementary Table 3:**
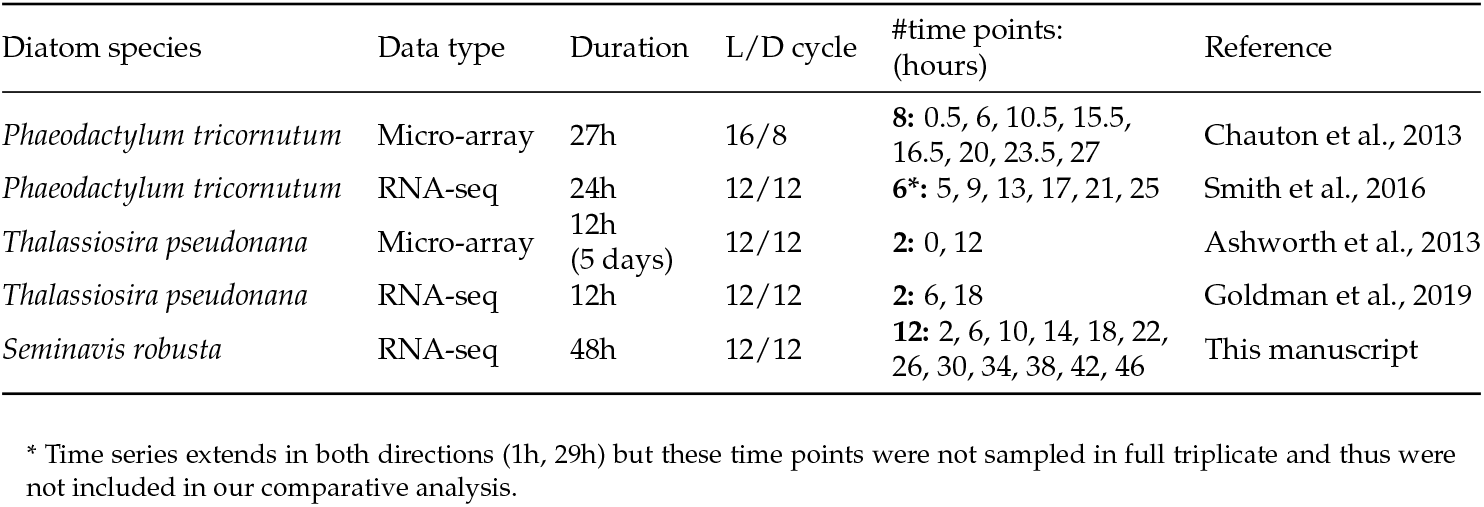
Overview of available day/night-related transcriptomes in diatoms. The “duration” column refers to the length of the experiment (time series), while “L/D cycle” indicates the photocycle regime to which cultures were entrained in each study. “#time points: (hours)” shows the number of time points sampled in each study (in bold), and the specific time points since the first illumination. The “main topic” column summarizes the main objective of each study. The study of Smith and colleagues (2016) was chosen for formal comparative analysis of the phasing of orthologous genes, as explained in the Methods.

| Diatom species | Data type | Duration | L/D cycle | #time points:<br>(hours) | Reference |
| --- | --- | --- | --- | --- | --- |
| *Phaeodactylum tricornutum* | Micro-array | 27h | 16/8 | **8:** 0.5, 6, 10.5, 15.5, 16.5, 20, 23.5, 27 | Chauton et al., 2013 |
| *Phaeodactylum tricornutum* | RNA-seq | 24h | 12/12 | **6\*:** 5, 9, 13, 17, 21, 25 | Smith et al., 2016 |
| *Thalassiosira pseudonana* | Micro-array | 12h<br>(5 days) | 12/12 | **2:** 0, 12 | Ashworth et al., 2013 |
| *Thalassiosira pseudonana* | RNA-seq | 12h | 12/12 | **2:** 6, 18 | Goldman et al., 2019 |
| *Seminavis robusta* | RNA-seq | 48h | 12/12 | **12:** 2, 6, 10, 14, 18, 22, 26, 30, 34, 38, 42, 46 | This manuscript |

\* Time series extends in both directions (1h, 29h) but these time points were not sampled in full triplicate and thus were not included in our comparative analysis.

